# Microstructural development from 9-14 years: evidence from the ABCD Study

**DOI:** 10.1101/2021.06.04.447102

**Authors:** Clare E Palmer, Diliana Pecheva, John R Iversen, Donald J Hagler, Leo Sugrue, Pierre Nedelec, Chun Chieh Fan, Wesley K Thompson, Terry L Jernigan, Anders M Dale

**Affiliations:** Center for Human Development, University of California, San Diego, 9500 Gilman Drive, La Jolla, CA 92161, USA; Center for Multimodal Imaging and Genetics, University of California, San Diego School of Medicine, 9444 Medical Center Dr, La Jolla, CA 92037, USA; Department of Radiology, University of California, San Diego School of Medicine, 9500 Gilman Drive, La Jolla, CA 92037, USA; Department of Radiology and Biomedical Imaging, University of California, San Francisco, 505 Parnassus Avenue, San Francisco, CA 94143, USA; Division of Biostatistics, Department of Family Medicine and Public Health, University of California, San Diego, La Jolla, CA, USA; Department of Cognitive Science, University of California, San Diego, 9500 Gilman Drive, La Jolla, CA 92161, USA; Department of Psychiatry, University of California, San Diego School of Medicine, 9500 Gilman Drive, La Jolla, CA 92037, USA; Department of Neuroscience, University of California, San Diego School of Medicine, 9500 Gilman Drive, La Jolla, CA 92037, USA

**Keywords:** development, neuroimaging, microstructure, subcortical, adolescence, diffusion

## Abstract

During late childhood behavioral changes, such as increased risk-taking and emotional reactivity, have been associated with the maturation of cortico-cortico and cortico-subcortical circuits. Understanding microstructural changes in both white matter and subcortical regions may aid our understanding of how individual differences in these behaviors emerge. Restriction spectrum imaging (RSI) is a framework for modelling diffusion-weighted imaging that decomposes the diffusion signal from a voxel into hindered, restricted, and free compartments. This yields greater specificity than conventional methods of characterizing diffusion. Using RSI, we quantified voxelwise restricted diffusion across the brain and measured age associations in a large sample (n=8,086) from the Adolescent Brain and Cognitive Development (ABCD) study aged 9-14 years. Older participants showed a higher restricted signal fraction across the brain, with the largest associations in subcortical regions, particularly the basal ganglia and ventral diencephalon. Importantly, age associations varied with respect to the cytoarchitecture within white matter fiber tracts and subcortical structures, for example age associations differed across thalamic nuclei. This suggests that age-related changes may map onto specific cell populations or circuits and highlights the utility of voxelwise compared to ROI-wise analyses. Future analyses will aim to understand the relevance of this microstructural developmental for behavioral outcomes.

## INTRODUCTION

Brain development during childhood and adolescence is associated with distributed structural alterations in both gray matter (GM) and white matter (WM) that occur concurrently with cognitive and behavioral development. WM tracts connect distributed neural networks across cortical and subcortical structures that are essential for a multitude of cognitive functions that continue to develop into late childhood (Baron Nelson et al., 2019; Peters et al., 2012; Simmonds et al., 2014). Alterations in reward and affective processing are particularly pertinent during adolescence (Casey et al., 2008) and are hypothesized to be underpinned by cortico-subcortical circuitry (Casey et al., 2016). The precise quantification of the microstructural changes during typical development may provide important information for understanding individual differences in cognition and the emergence of increased emotional reactivity and risk-taking in this period. Diffusion tensor imaging (DTI) has frequently been used to probe microstructural changes in the brain. Previous studies have shown increases in fractional anisotropy (FA) and decreases in mean diffusivity (MD) throughout the brain across childhood and into young adulthood, with variability in the trajectory of microstructural development across different brain regions (for review see Lebel & Deoni, 2018). Many studies have measured developmental changes in DTI metrics within WM (Krogsrud et al., 2016a; Catherine Lebel & Beaulieu, 2011; Pohl et al., 2016), but fewer studies have explored DTI changes in deep gray matter structures, in part due to the inadequacies of DTI for studying complex cytoarchitecture and the lower signal-to-noise ratio (SNR) when estimating FA in particular (Farrell et al., 2007). Despite this, in one study, increases in FA from 5-30 years appeared to be larger in subcortical regions compared to the WM tracts (Lebel et al., 2008).

The diffusion tensor model only allows the expression of a single principal direction of diffusion and is unable to adequately represent mixtures of neurite orientations within a voxel. Recent advances in diffusion data acquisition, including multiple b-value acquisitions and high angular resolution diffusion imaging (HARDI), have enabled more complex models of tissue microstructure, taking into account multiple tissue compartments, multiple fiber populations in WM and orientated structure of neurites (axons and dendrites) within GM and WM. Restriction spectrum imaging (RSI; (Brunsing et al., 2017; White et al., 2013; White et al., 2013, 2014) uses multiple b-value HARDI data to model the diffusion-weighted signal as emanating from multiple tissue compartments, reflecting free, hindered and restricted water, with different intrinsic diffusion properties. The hindered compartment is thought to primarily represent extracellular space although may also describe diffusion within intracellular spaces with dimensions larger than the diffusion length scale (typically, ∼10μm, for the diffusion sequences used in human imaging studies). The restricted compartment is thought to primarily represent intracellular space, within cells or processes of dimensions smaller than the diffusion length scale. Free water diffusion primarily represents cerebrospinal fluid (CSF) or intravascular spaces. Within each voxel, RSI models the diffusion signal as a linear mixture of these different compartments. Spherical deconvolution (SD) is used to reconstruct the fiber orientation distribution (FOD) in each voxel for each compartment.

Using RSI, we can quantify the relative proportion of restricted, hindered and free water diffusion within each voxel of the brain. The signal fraction for each compartment is normalized by total diffusion signal across all compartments (restricted normalized total signal fraction, RNT; hindered normalized total signal fraction, HNT; free normalized total signal fraction, FNT^1^). Moreover, from the spherical harmonic coefficients (SH) from the RSI model, we can estimate the signal fraction of restricted normalized directional (anisotropic) diffusion (RND) and restricted normalized isotropic diffusion (RNI). By dividing RND by RNT we can additionally estimate the relative proportion of directional to isotropic diffusion specifically within the restricted compartment (restricted directional fraction (RDF), and how this changes with age. There are several developmental processes that can modulate the relative proportion of restricted to hindered diffusion within a voxel (see Table 1). For example, myelination increases RNT relative to HNT by both decreasing the extracellular space and decreasing the exchange of water molecules across the axonal membrane. Dendritic sprouting, arborization and increases in neurite density can also increase the RNT by decreasing the proportion of extracellular space within a voxel. The relative size and shape of restricted compartments will then differentially modulate isotropic and anisotropic diffusion.

**Table 1.**
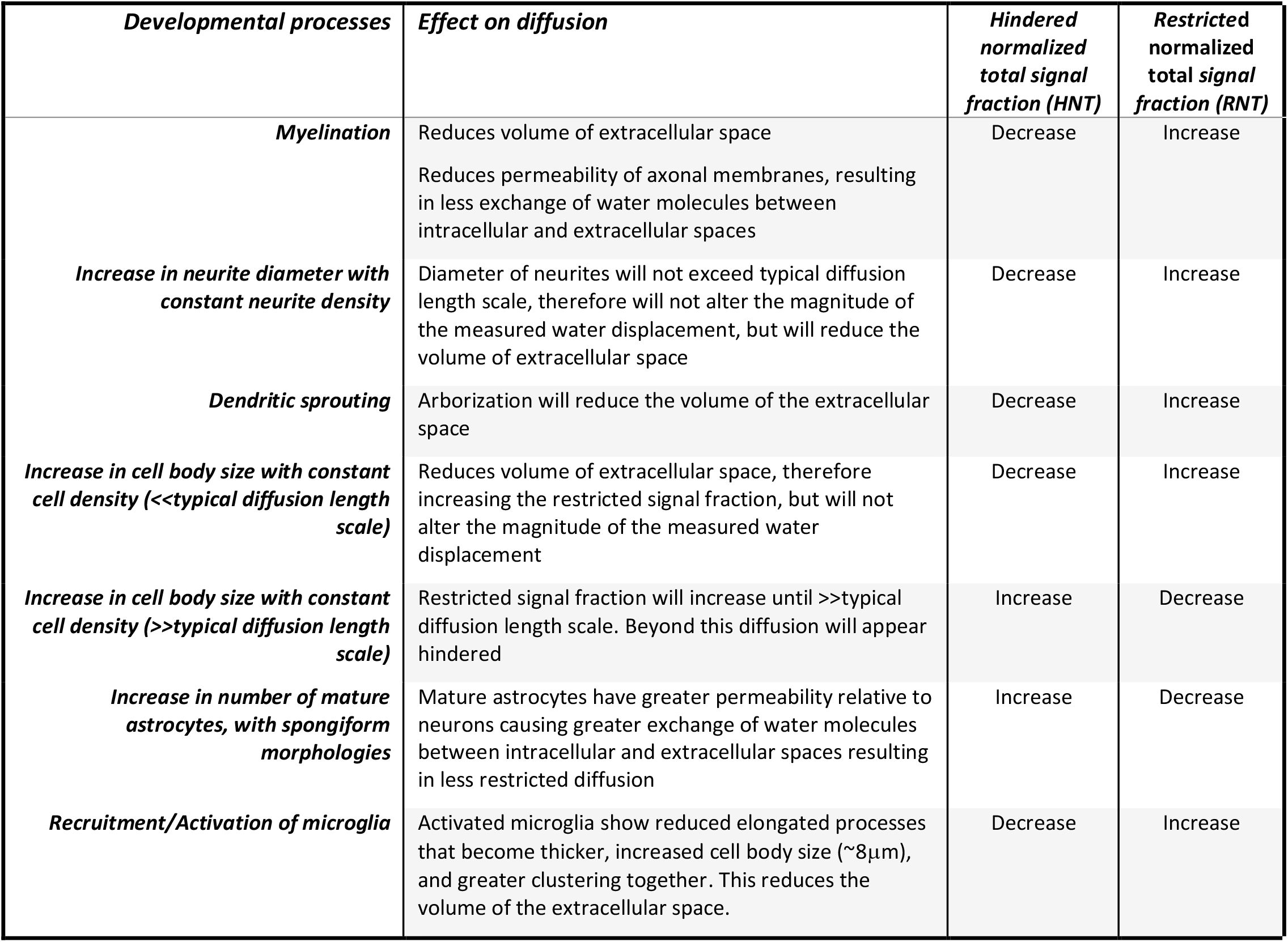
Outline of how different developmental cellular processes can modulate both the hindered and restricted signal fractions.

RSI has been used in several different applications (Carper et al., 2017; Loi et al., 2016; Reas et al., 2017, 2020), but has not previously been used to study developmental changes in late childhood. However, similar multi-compartment models, such as neurite orientation dispersion and density imaging (NODDI), have been shown to be more sensitive to developmental changes than DTI metrics (Genc, Malpas, et al., 2017). Neurite density index (NDI), from the NODDI model, which reflects the intracellular volume fraction, was positively associated with age across all WM tracts in several recent studies (Geeraert et al., 2019; Genc, Malpas, et al., 2017; Lynch et al., 2020; Mah et al., 2017). However, the orientation dispersion index (ODI), a measure of the degree of dispersion of intracellular diffusion, showed no age associations, suggesting that WM development across childhood and adolescence is not associated with changes in neurite coherence (Genc, Malpas, et al., 2017; Lynch et al., 2020). Increases in the intracellular volume fraction from the NODDI model have also been shown to significantly increase with age from 8-13 years in subcortical regions. These results suggest that age related increases in restricted diffusion measured with more sensitive multi-compartment models are apparent in both WM and deep GM. Although NODDI is a useful model for describing intracellular diffusion, NDI is limited in that it represents a measure of the total intracellular volume fraction; in contrast, RSI can delineate isotropic and anisotropic diffusion within the restricted compartment. For example, in voxels with crossing fibers that are oriented perpendicular to one another, ODI will be estimated to be very high; whereas RND will provide a more accurate estimation of the anisotropy of the two coherent crossing fibers. RSI therefore provides differential information about intracellular diffusion within each voxel compared to previously explored multi-compartment models.

In the current study, we have used longitudinal data across two time points to estimate age-related changes in tissue microstructure in WM and subcortical regions. We have used data from Release 4.0 of the Adolescent Brain and Cognitive Development (ABCD) Study to measure whole-brain voxelwise age associations with the total restricted, hindered and free water signal fractions, as well as the restricted isotropic and anisotropic signal fractions form the RSI model. The large sample size (n=8086) and small age range at each time point (9-11 years at baseline; 11-14 years at follow-up) provides high precision to delineate microstructural changes with age across the brain.

## METHODS

### Sample

The ABCD study is a longitudinal study across 21 data acquisition sites following 11,880 children starting at 9-11 years. This paper uses baseline and two-year follow up (FU) data from the NIMH Data Archive ABCD Collection Release 4.0 (DOI: 10.15154/1523041). The ABCD cohort is epidemiologically informed (Garavan et al., 2018), including participants from demographically diverse backgrounds, and has an embedded twin cohort and many siblings. Exclusion criteria for participation in the ABCD Study were limited to: 1) lack of English proficiency in the child; 2) the presence of severe sensory, neurological, medical or intellectual limitations that would inhibit the child’s ability to comply with the protocol; 3) an inability to complete an MRI scan at baseline. The study protocols are approved by the University of California, San Diego Institutional Review Board. Parent/caregiver permission and child assent were obtained from each participant.

All statistical analyses included 14,043 observations with 8,086 unique subjects, such that 5,957 participants had data at two time points. Participants were aged from 107-166 months (8.9-13.8 years). Observations were included in the final sample if the participant had complete data across sociodemographic factors (household income, highest parental education, ethnicity), available genetic data (to provide ancestry information using the top 10 principal components), available imaging data that passed all inclusion criteria and available information regarding acquisition scanner ID and software version. In the ABCD Study, Release 4.0, there are 19,658 available scans with scanner information (12% missingness). Of these scans, 2,655 were excluded for not meeting the recommended imaging inclusion criteria outlined in the Release 4.0 release notes and supplementary table 1 (imaging scans were included if: imgincl_dmri_include==1 & imgincl_t1w_include==1 & mrif_score<3), and an additional 90 observations were excluded for poor registration defined below (in *Atlas Registration*). The final sample included all remaining observations that had complete data for the previously listed information. Table 2 shows the demographics of the final sample used for statistical analysis stratified by time-point. Participants who had completed their 2 year FU in Release 4.0 were more likely to have higher household income and have male assigned as their sex at birth. This may reflect differences in recruitment procedures over the course of recruitment in order to ensure the final sample reflected the demographics of the US population as closely as possible.

**Table 2.**
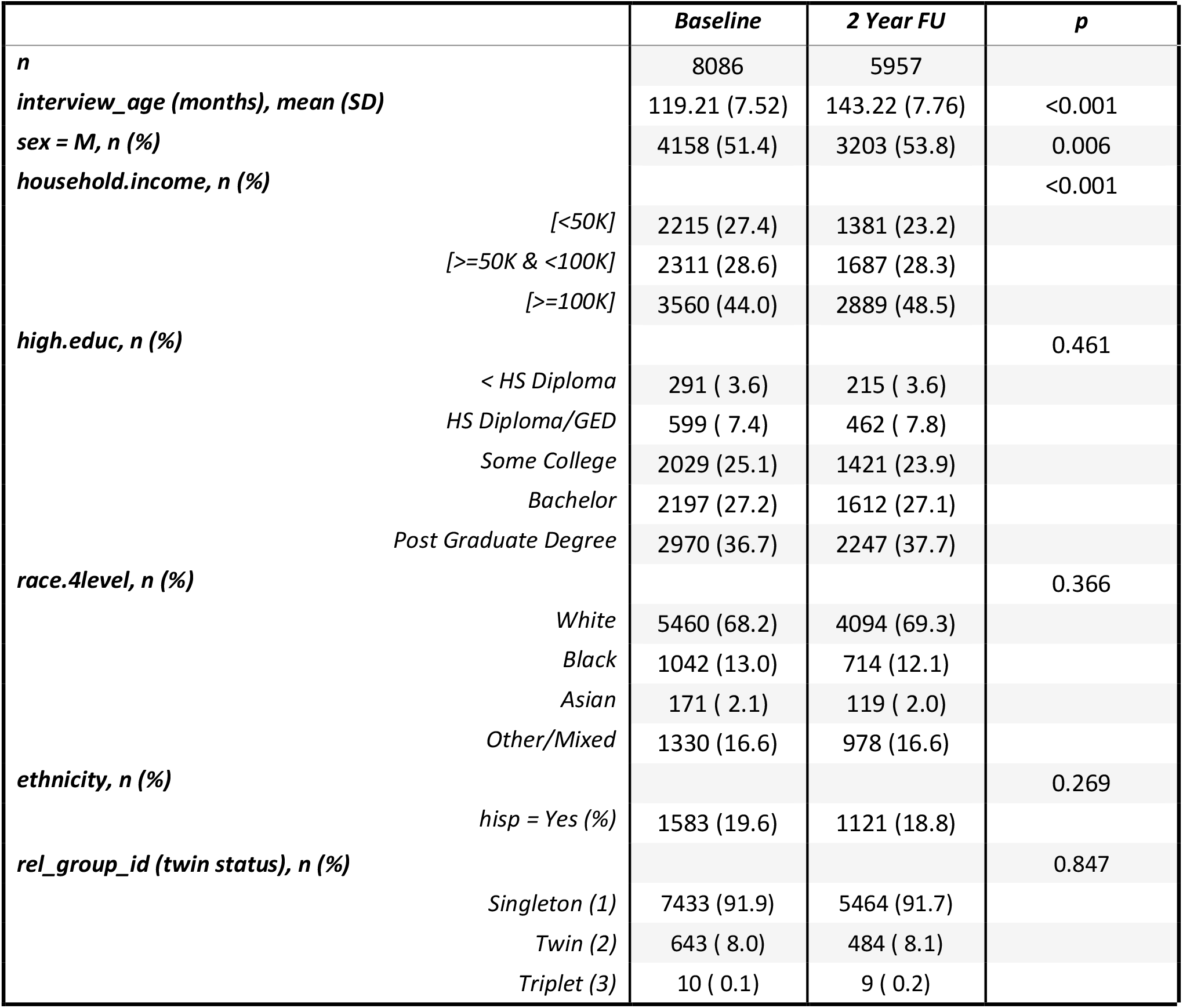
Demographics of the sample. Demographic data is shown for age in months (mean, (SD)), sex at birth, household income, parental education, self-declared race, endorsement of Hispanic ethnicity and self-declared twin/triplet status (n, (%)). These factors are stratified by time point: baseline and 2-year FU. There were significant differences in income and sex at birth for those who had 2-year FU data in Release 4.0 indicative of differences in the demographics of participants as they were recruited. Participants recruited earlier in the study were more likely to have higher household income and be born male. All of these variables are controlled for in all statistical analyses to account for this. Variable names from the tabulated data release are included in the table for replication.

### MRI acquisition

The ABCD MRI data were collected across 21 research sites using Siemens Prisma, GE 750 and Philips Achieva and Ingenia 3T scanners. Scanning protocols were harmonized across sites. Full details of structural and diffusion imaging acquisition protocols used in the ABCD study have been described previously (Casey et al., 2018; Hagler et al., 2019) so only a short overview is given here. dMRI data were acquired in the axial plane at 1.7mm isotropic resolution with multiband acceleration factor 3. Diffusion-weighted images were collected with seven b=0 s/mm^2^ frames and 96 non-collinear gradient directions, with 6 directions at b=500 s/mm^2^, 15 directions at b=1000 s/mm^2^, 15 directions at b=2000 s/mm^2^, and 60 directions at b=3000 s/mm^2^. T1-weighted images were acquired using a 3D magnetization-prepared rapid acquisition gradient echo (MPRAGE) scan with 1mm isotropic resolution and no multiband acceleration. 3D T2-weighted fast spin echo with variable flip angle scans were acquired at 1mm isotropic resolution with no multiband acceleration.

### Image Processing

The processing steps for diffusion and structural MR data are outlined in detail in Hagler et al., (2019). Briefly, dMRI data were corrected for eddy current distortion using a diffusion gradient model-based approach (Zhuang et al., 2006). To correct images for head motion, we rigid-body-registered each frame to the corresponding volume synthesized from a robust tensor fit, accounting for image contrast variation between frames. Dark slices caused by abrupt head motion were replaced with values synthesized from the robust tensor fit, and the diffusion gradient matrix was adjusted for head rotation (Hagler et al., 2009, 2019). Spatial and intensity distortions caused by B0 field inhomogeneity were corrected using FSL’s *topup* (Andersson et al., 2003) and gradient nonlinearity distortions were corrected for each frame (Jovicich et al., 2006). The dMRI data were registered to T1w structural images using mutual information (Wells et al., 1996) after coarse pre-alignment via within-modality registration to atlas brains. dMRI data were then resampled to 1.7 mm isotropic resolution, equal to the dMRI acquisition resolution.

T1w and T2w structural images were corrected for gradient nonlinearity distortions using scanner-specific, nonlinear transformations provided by MRI scanner manufacturers (Jovicich et al., 2006; Wald et al., 2001) and T2w images are registered to T1w images using mutual information (Wells et al., 1996). Intensity inhomogeneity correction was performed by applying smoothly varying, estimated B1-bias field (Hagler et al., 2019). Images were rigidly registered and resampled into alignment with a pre-existing, in-house, averaged, reference brain with 1.0 mm isotropic resolution (Hagler et al., 2019).

### Microstructural models

#### Restriction spectrum imaging (RSI)

The RSI model was fit to the diffusion data to model the diffusion properties of the cerebral tissue (Nathan S. White et al., 2013, 2014). RSI estimates the relative fraction that separable pools of water within a tissue contribute to the diffusion signal, based on their intrinsic diffusion characteristics. Free water (e.g., CSF) is defined by unimpeded water diffusion. Hindered diffusion follows a Gaussian displacement pattern characterised by the presence of neurites, glia and other cells. This includes water both within the extracellular matrix and certain intracellular spaces with dimensions larger than the diffusion length scale (typically, ∼10μm, for the diffusion sequences used in human imaging studies (Nathan S. White et al., 2013)). Restricted diffusion describes water within intracellular spaces confined by cell membranes and follows a non-Gaussian pattern of displacement. Imaging scan parameters determine the sensitivity of the diffusion signal to diffusion within these separable pools. At intermediate b-values (b=500-2500s/mm^2^), the signal is sensitive to both hindered and restricted diffusion; whereas, at high b-values (b≥3000s/mm^2^), the signal is primarily sensitive to restricted diffusion. The hindered and restricted compartments are modeled as fourth order spherical harmonic (SH) functions and the free water compartment is modelled using zeroth order SH functions. The axial diffusivity (AD) is held constant, with a value of 1 × 10^−3^ mm^2^/s for the restricted and hindered compartments. For the restricted compartment, the radial diffusivity (RD) is fixed to 0 mm^2^/s. For the hindered compartment, RD is fixed to 0.9 × 10^−3^ mm^2^/s. For the free water compartment the isotropic diffusivity is fixed to 3 × 10^−3^ mm^2^/s. Theoretically, any increases in the tortuosity of the hindered compartment, for example due to a decrease in the volume of the extracellular space, will decrease the effective diffusivity in the hindered compartment; however, in our model we are assuming the hindered diffusivity is constant. Spherical deconvolution (SD) is used to reconstruct the fiber orientation distribution (FOD) in each voxel from the restricted compartment. The restricted directional measure, RND, is the norm of the SH coefficients for the second and fourth order SH coefficients (divided by the norm of all the coefficients across the restricted, hindered and free water compartments). This models oriented diffusion emanating from multiple directions within a voxel.

The restricted isotropic measure, RNI, refers to the spherical mean of the FOD across all orientations (zeroth order SH divided by the norm of all the coefficients across the restricted, hindered and free water compartments). The sum of these measures is the restricted normalized total signal fraction, RNT.

In this study we explore associations between age and the rotation-invariant features of the restricted compartment FOD. For a detailed description of the derivation of the RSI model see (N.S. White et al., 2013; Nathan S. White et al., 2013). We extracted a measure of the restricted isotropic and restricted anisotropic diffusion signal. Within each voxel the total diffusion signal, S, can be represented as

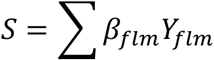

where *Y*_*flm*_ is a SH basis function of order *l* and degree *m* of the FOD corresponding to the *f*th compartment, and *β*_*flm*_ are the corresponding SH coefficients. The total restricted normalized signal fraction (RNT), normalized by all compartments, is defined as follows:

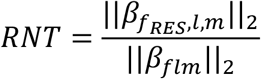

The total hindered normalized signal fraction (HNT), normalized by all compartments, is defined as follows:

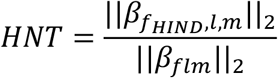

The total free water normalized signal fraction (FNT), normalized by all compartments, is defined as follows:

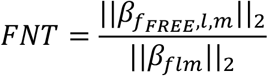

The measure of the restricted normalized isotropic signal fraction is given by the coefficient of the zeroth order SH coefficient, 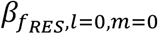, where *f*_*RES*_ is the restricted compartment, normalized by the Euclidian norm of all *β*_*flm*_ and termed RNI:

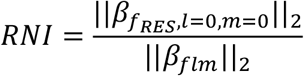

The measure of the restricted normalized directional signal fraction is given by the norm of 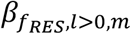, where *l* > 0, and *f*_*RES*_ is the restricted compartment, and is termed RND:

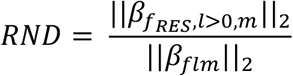

These normalized RSI measures are unitless and range from 0 to 1. Given that RNI and RND are both normalized by the SH coefficient across all compartments, changes in the overall restricted or hindered signal fractions can modulate both of these measures similarly. To determine the relative contribution of isotropic to anisotropic diffusion solely within the restricted compartment we estimated the proportion of RND over RNT, termed RDF.

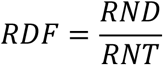

The magnitude of diffusion that we are sensitive to is dependent on the diffusion scan parameters. Typical diffusion times used in clinical DWI scans are approximately 10–50ms corresponding to average molecular displacements on the order of 10μm (Mukherjee et al., 2008). Any water displacements smaller than this scale would not result in detectable dephasing, regardless of b-value, therefore would not lead to changes in the measured diffusion coefficient. However, changes in cell size <∼10μm could alter the relative signal fractions of hindered and restricted diffusion in a voxel. Diffusion estimated in these compartments is also dependent on the permeability of cellular membranes; greater exchange across intracellular and extracellular space will mean that diffusion will appear more hindered rather than restricted. Table 1 outlines the expected changes to the hindered and restricted signal fractions following example microstructural developmental processes.

#### Diffusion tensor imaging

The diffusion tensor model (Basser et al., 1994; Basser & Pierpaoli, 1996) was used to calculate fractional anisotropy (FA) and mean diffusivity (MD). Diffusion tensor parameters were calculated using a standard, linear estimation approach with log-transformed diffusion-weighted (DW) signals (Basser et al., 1994). Tensor matrices were diagonalized using singular value decomposition, obtaining three eigenvectors and three corresponding eigenvalues. FA and MD were calculated from the eigenvalues (Basser & Pierpaoli, 1996).

### Atlas registration

To allow for voxelwise analysis, subjects’ imaging data were aligned using a multimodal nonlinear elastic registration algorithm. At the end of the preprocessing steps outlined in *Image Processing* and described in detail in Hagler et al. (2019), subjects’ structural images and diffusion parameter maps were aligned to an ABCD-specific atlas, using a custom diffeomorphic registration method (Holland & Dale, 2011). The ABCD-specific atlas was constructed from n=17,636 ABCD participants aged 9-14 years using an iterative procedure, consisting of an initial affine registration, followed by a multi-scale, multi-channel elastic diffeomorphic registration. Eleven input channels were used for the multimodal registration: 3D T1, zeroth and second order SH coefficients from the restricted FOD, zeroth order SH coefficient from the hindered and free water FODs, white matter and grey matter segmentations. After each iteration, morphed volumes for each subject were averaged to create an updated atlas, and then the process was repeated until convergence. Participants with poor registration to atlas were excluded from the average and subsequent statistical analyses. Poor registration was defined as a mean voxelwise correlation to atlas across channels <0.8 (see *Sample* for number excluded).

### Labelling regions of interest (ROI)

Major white matter tracts were labelled using AtlasTrack, a probabilistic atlas-based method for automated segmentation of white matter fiber tracts (Hagler et al., 2009, 2019). Unilateral binary masks for each ROI (except the CC, Fmaj and Fmin which are interhemispheric) were created by thresholding at 0.9 probability across the ROI meaning that in a given voxel at least 90% of participants showed that ROI label. A list of WM tract ROIs used in this study is listed in Supplementary Table 2. Subcortical structures were labeled using the Freesurfer 5.3 segmentation (Fischl et al., 2002). Subjects’ native space Freesurfer parcellations were warped to the atlas space and averaged across subjects. Bilateral binary masks for each ROI were created using a probabilistic threshold of 0.9 with the same meaning as above. Additional subcortical nuclei, not available in the FreeSurfer segmentation, were labeled by registering readily available, downloadable, high spatial resolution atlases to our atlas space. The Pauli atlas was generated using T1 and T2 scans from 168 typical adults from the Human Connectome Project (HCP) (Pauli et al., 2018). The Najdenovska thalamic nuclei atlas was generated using a k-means algorithm taking as inputs the mean FOD SH coefficients from within a Freesurfer parcellation of the thalamus, using adult HCP data from 70 subjects (Najdenovska et al., 2018). Bilateral binary masks were created for all ROIs in atlas space. All subcortical ROIs and abbreviations are listed in Supplementary Table 3.

### Statistical analysis

#### Voxelwise analyses

Univariate general linear mixed effects models (GLMMs) were applied at each voxel to test the associations between age and diffusion metrics (RNT, HNT, FNT, RNI, RND, RDF, FA and MD) as the dependent variables. All of the main results shown are from a linear model (model below) with age included as a single predictor in long format and the longitudinal component modelled as a random effect of subject. Results were also compared against a model with an age*sex interaction and are reported in the supplementary analyses. Given the genetic relatedness within the sample, family relatedness was also controlled for as a random effect. Given the demographic diversity in the sample, all statistical analyses controlled for the sociodemographic variables household income, parental education and Hispanic ethnicity and the top 10 genetic principal components were used to account for ancestry effects in lieu of self-declared race. Additional fixed effects included scanner ID, MRI software version and motion (average frame-wise displacement in mm).

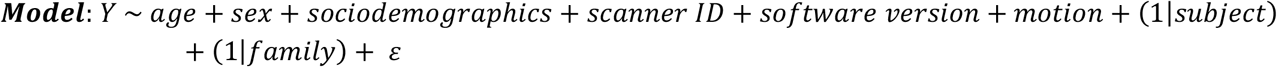

Whole-brain voxelwise analyses were corrected for multiple comparisons at an alpha level of 0.05 using a Bonferroni correction across 156,662 voxels to provide a voxelwise corrected threshold of p = 0.05 / 156,662 = 3.19e-7, corresponding to |t| = 4.98. This provides a conservative estimate of significant developmental effects as the true number of independent tests is likely to be smaller than this. Unthresholded t-statistic maps are presented in the main figures with the Bonferroni significance threshold marked on the colorbar. This provides a comprehensive description of the continuous distribution of effects beyond this conservative boundary. All imaging metrics were rank normalized prior to statistical analysis to adhere to normality assumptions of the linear model.

#### Region-of-interest (ROI) Analyses

ROI analyses were also conducted using the same GLMM. The dependent variable for each ROI for each diffusion metric was calculated by taking the mean diffusion metric across the voxels within each ROI mask. Violin plots were generated to show the variability in voxelwise effects across all voxels within each ROI mask in order to highlight the heterogeneity of developmental effects within each ROI. ROI analyses were corrected for multiple comparisons at an alpha level of 0.05 using a Bonferroni correction across 49 ROIs to provide a voxelwise corrected threshold of p = 0.05 / 49 = 0.0010, corresponding to |t| = 3.08. All ROIs were rank normalized prior to statistical analysis to adhere to normality assumptions of the linear model.

All statistical analyses were conducted using custom code in MATLAB v2017a. Code will be available on GITHUB.

#### Estimation of scanner and software version effects

Scanner and software effects were estimated across the brain by estimating a mean voxelwise change in pseudo-R^2^ (Δ*pseudoR*^2^) from a full model (all predictors) to a reduced model (either no dummy-coded scanner or software version predictors).

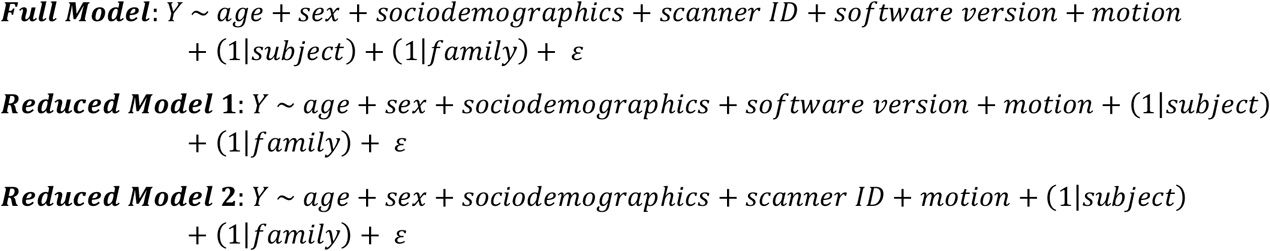

Pseudo-*R*^2^ was calculated using the below equation where *Ŷ* is a matrix of the voxelwise predicted imaging values for a given modality generated by the full or reduced model, and *Y* is the matrix of voxelwise observed imaging values. The variance of *Ŷ* and *Y* (across participants) was averaged across voxels before dividing to produce the pseudo-*R*^2^ estimate as a mean estimate across voxels.

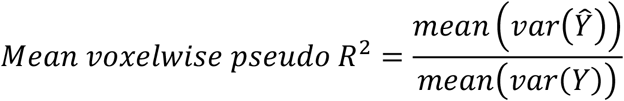

Δ*pseudoR*^2^ was calculated by taking the difference between the pseudo-*R*^2^ estimates for the full and reduced models.

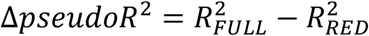

The variance in each imaging metric explained by scanner and software version predictors (as defined using Δ*pseudoR*^2^) is outlined in supplementary table 4.

## RESULTS

### Mean voxelwise RSI metrics across participants

The RSI model estimates diffusion within different compartments and these metrics are normalized into signal fractions in order to determine the relative proportion of restricted, hindered and free water diffusion within each voxel. Figure 1A-F shows mean voxelwise maps across participants of these normalized signal fractions. The restricted normalized total signal fraction (RNT) was largest within the WM and lowest within the GM. In contrast, the hindered normalized total signal fraction (HNT) was largest within the GM and lowest within the WM. The free water normalized total signal fraction (FNT) was low in brain tissue and high within the CSF. These normalized metrics sum to 1, therefore increases in the relative signal fraction of one of these compartments with age will result in decreases in at least one other compartment. Within the restricted compartment specifically we have separated the proportion of isotropic and directional diffusion into dissociable metrics (figure 1G-L). The restricted normalized directional signal fraction (RND) shows a much greater contrast between WM and GM (with higher values in WM) compared to the restricted normalized isotropic signal fraction (RNI). Increases in RNT with age will lead to increases in both RNI and RND; however, the specific microstructural changes occurring can lead to differential changes in RNI compared to RND. We have calculated the proportion of restricted directional over the total restricted diffusion (RDF) in order to determine changes in the relative proportion of isotropic to anisotropic diffusion with age. Mean values of RDF were greater within WM voxels compared to GM voxels reflecting the greater contrast in RNI vs RND in WM. These maps can be compared to T1-weighted images shown in Figure 1M,N.

**Figure 1.**
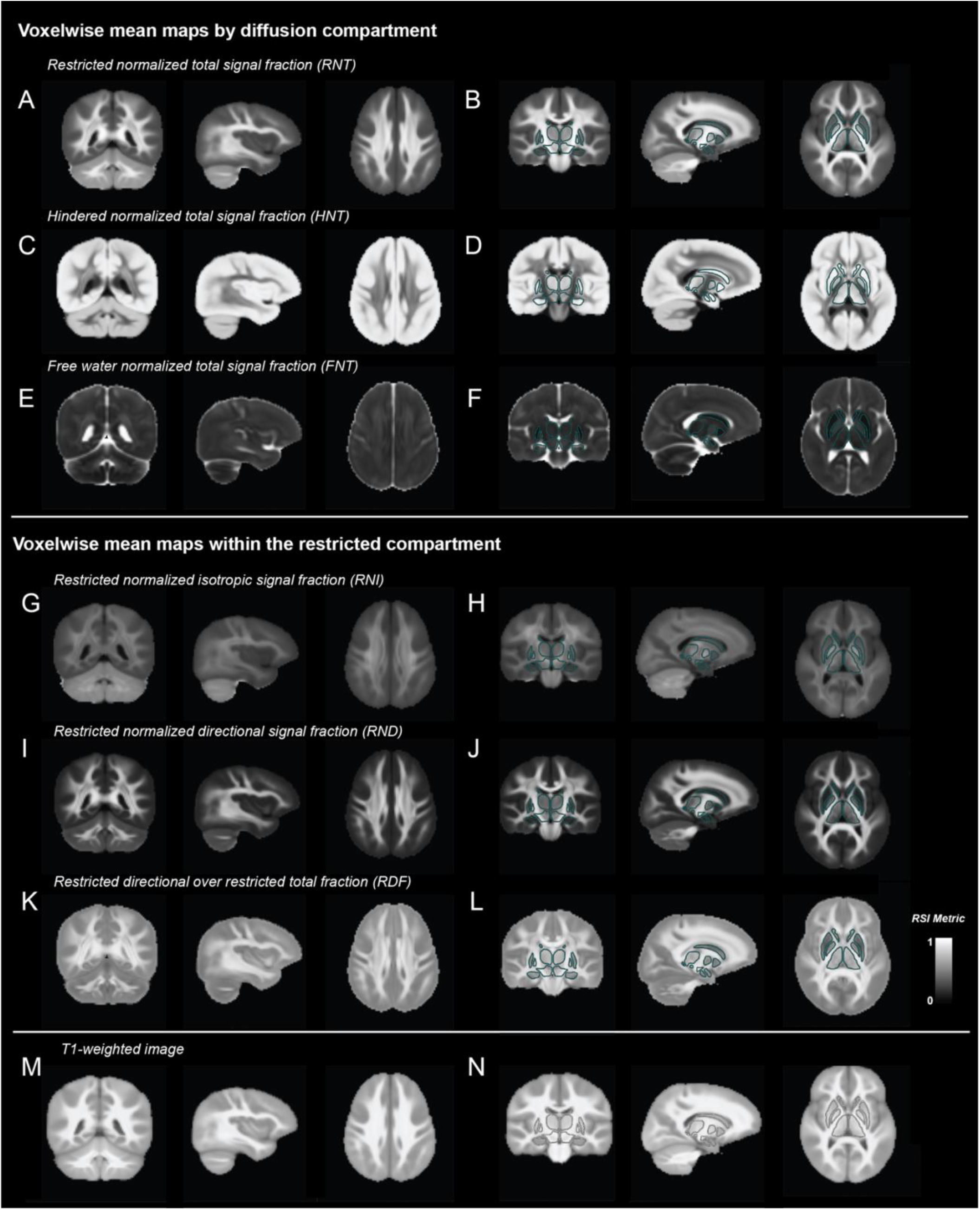
Voxelwise mean maps of diffusion metrics and T1-weighted images. Top panel: voxelwise mean maps for RSI metrics across different compartments: RNT (A,B), HNT (C,D) and FNT (E,F). Middle panel: voxelwise mean maps for RSI metrics within the restricted compartment: RNI (G,H), RND (I,J) and RDF (K,L). Bottom panel: voxelwise mean T1-weighted images (M,N).

**Figure 2.**
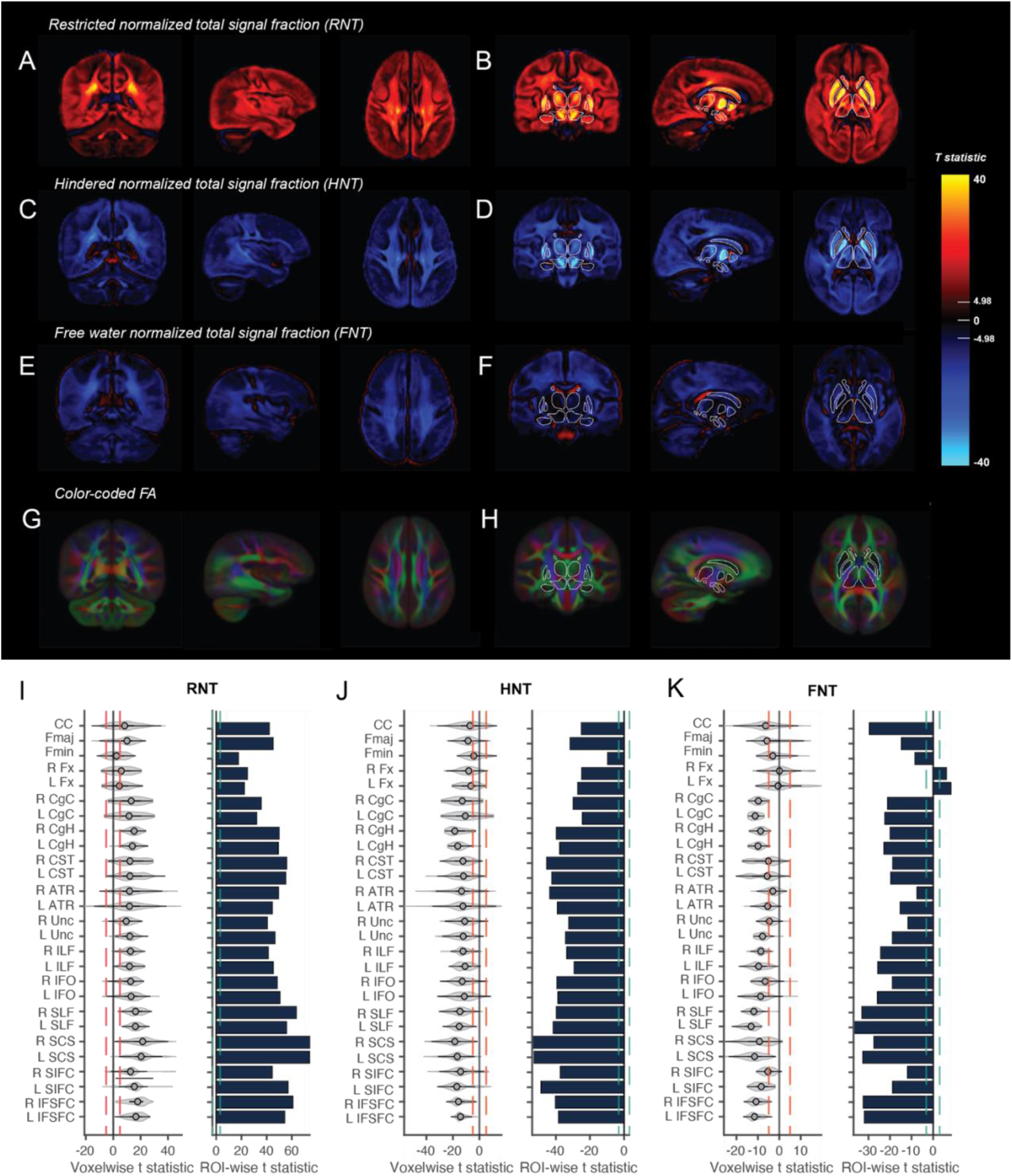
Associations between age and RSI compartment signal fractions. Voxelwise t-statistics for the association between age and RNT (A,B), HNT (C,D) and FNT (E,F) across different brain slices. Effects are unthresholded. Voxelwise Bonferroni corrected significance threshold (|t|=4.98) is marked on the colorbar. Outlines of the subcortical FreeSurfer ROIs are overlaid for the thalamus, caudate, pallidum, putamen, ventral diencephalon, amygdala and hippocampus to orient the reader. G,H) Color coded FA showing the primary diffusion direction in each voxel from the tensor model. Larger versions of the same slices with WM fiber tracts labeled are shown in supplementary figure 1. I-K) Violin plots show the distribution of voxelwise t-statistics extracted from each WM fiber tract. Red dotted lines show voxelwise Bonferroni corrected significance threshold. Bar plots show t-statistics from ROI analyses for the mean RSI metrics from each WM fiber tract. Green dotted line shows ROI Bonferroni corrected significance threshold (|t|=3.08). Plots are shown for RNT (G), HNT (H) and FNT (I). WM tract ROI abbreviations described in Supplementary Table 2.

### Age associations in WM across the different RSI compartments

Voxelwise associations between age and RNT were positive across the brain, such that the proportion of the diffusion signal within each voxel that was restricted increased with age (Figure 3A,B). The largest voxelwise effects were found in subcortical regions. Inverse associations were found for the hindered signal fraction, as expected given the normalization across these metrics (Figure 3C,D). Voxelwise age associations with FNT were also negative across GM and WM voxels, positive within the ventricles and limited with subcortical structures (Figure 3E,F). However, it is important to note that the proportion of free water within the brain tissue is very small as highlighted in the mean voxelwise FNT maps (Fig 1E,F), therefore although these age associations are significant across subjects the relative magnitude of the effect compared to changes in RNT and HNT is very low. There were no age associations with FNT within the deep GM structures.

**Figure 3.**
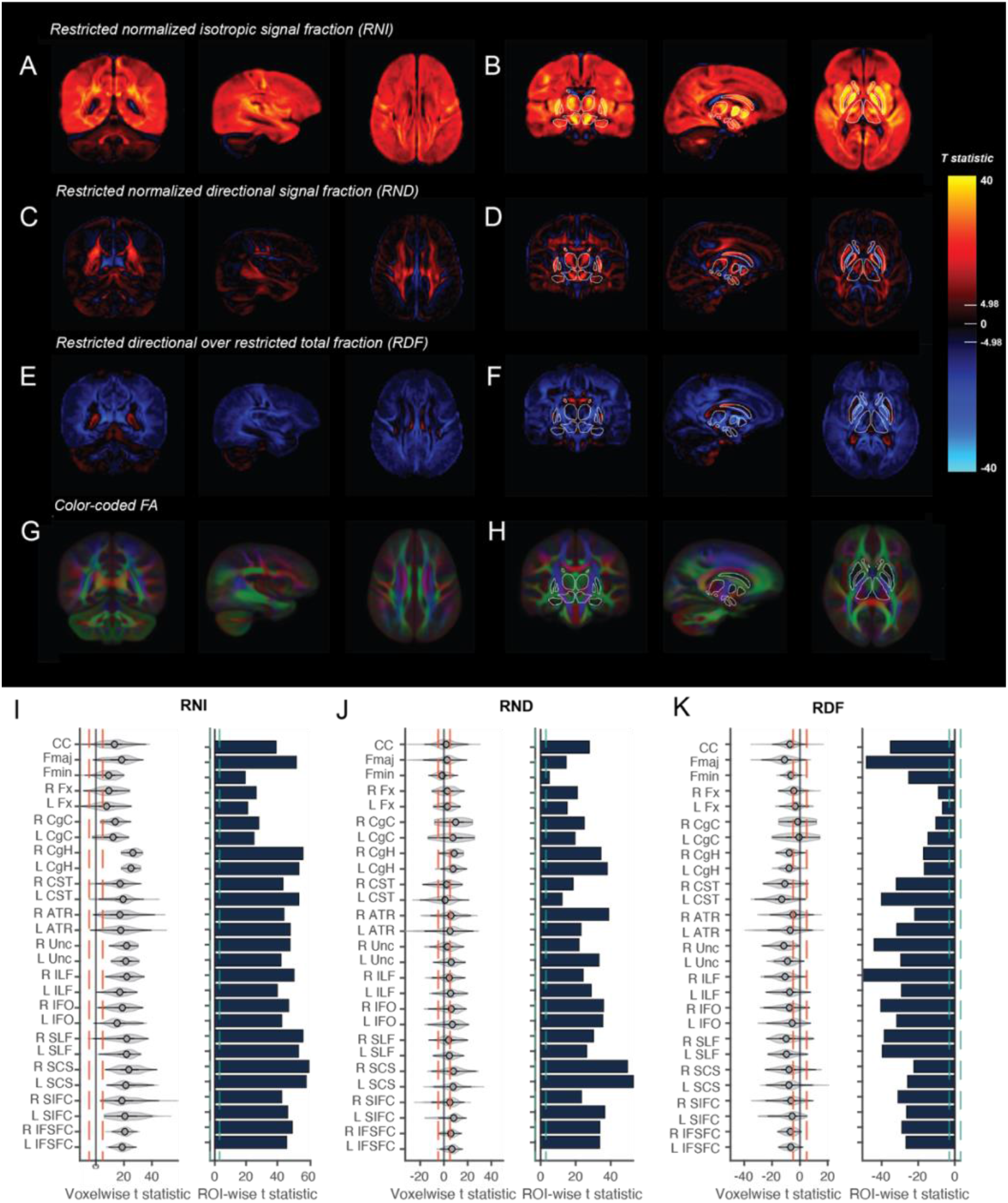
Associations between age and RSI metrics within the restricted compartment. Voxelwise t-statistics for the association between age and RNI (A,B), RND (C,D) and RDF (E,F) across different brain slices. Effects are unthresholded. Voxelwise Bonferroni corrected significance threshold (|t|=4.98) is marked on the colorbar. Outlines of the subcortical FreeSurfer ROIs are overlaid for the thalamus, caudate, pallidum, putamen, ventral diencephalon, amygdala and hippocampus to orient the reader. G,H) Color coded FA showing the primary diffusion direction in each voxel from the tensor model. Larger versions of the same slices with WM fiber tracts labeled are shown in supplementary figure 1. I-K) Violin plots show the distribution of voxelwise t-statistics extracted from each WM fiber tract. Red dotted lines show voxelwise Bonferroni corrected significance threshold. Bar plots show t-statistics from ROI analyses for the mean RSI metrics from each WM fiber tract. Green dotted line shows ROI Bonferroni corrected significance threshold (|t|=3.08). Plots are shown for RNI (G), RND (H) and RDF (I). WM tract ROI abbreviations are outlined in Supplementary Table 2.

A probabilistic atlas-based method for automated segmentation was used to determine ROIs for the major WM fiber tracts. Color coded voxelwise FA maps highlight the primary direction of diffusion across the brain (Fig 3G,H) and enable comparison of where the main WM fiber tracts are located. Larger versions of the same maps with WM fiber tracts labeled are shown in supplementary figure 1. Voxelwise age associations were extracted for each WM fiber tract in order to determine the distribution of effects within these ROIs. Violin plots highlight the heterogeneity of age-related changes in these RSI metrics within the WM fibers (Figure 3I-K). Moreover, these figures demonstrate the proportion of voxels above and below the conservative Bonferroni corrected threshold for whole-brain voxelwise statistical significance (red dotted line). ROI analyses were also conducted on the mean RSI metric within each fiber tract. For RNT and HNT, all WM fiber tracts showed highly significant age associations indicative of a global increase in the restricted signal fraction with age. Interestingly, WM tracts with voxels near to or innervating subcortical regions appeared to show the greatest heterogeneity and largest age associations, for example the anterior thalamic radiations (ATR), the corticospinal tract (CST), which innervates the ventral diencephalon (VDC), the superior cortico-striate (SCS) and the striatal inferior frontal cortex (SIFC) tract. The forceps minor (Fmin) and bilateral fornix (Fnx) showed the smallest age associations across the RSI compartments. Supplementary tables 5-7 show summary statistics for the voxelwise and ROIwise analyses within each WM fiber tract for RNT, HNT and FNT.

There were no significant voxelwise age-by-sex interaction effects for RNT, HNT or FNT at the Bonferroni corrected significance threshold. Voxelwise age associations in a model without an age-by-sex interaction were highly correlated with a model including the interaction term (supplementary figure 2A-C). All main effects presented are from models without an age-by-sex interaction.

### Age associations in WM within the restricted compartment

Voxelwise associations between age and both RNI and RND were positive across the brain reflecting the increase in the total restricted signal fraction. Voxelwise age associations with RNI were widespread across the brain and largest in deep GM structures (Figure 4A,B). Voxelwise associations with RND were smaller than RNI and more concentrated along the center of the main WM tracts (Figure 4C,D). Voxelwise associations with RDF (the fraction of restricted directional diffusion over RNT) were negative across the WM highlighting that the relative proportion of directional to isotropic diffusion within the restricted compartment decreased with age i.e. the proportion of isotropic restricted diffusion increased at a greater rate compared to the proportion of directional diffusion (Figure 4E,F).

**Figure 4.**
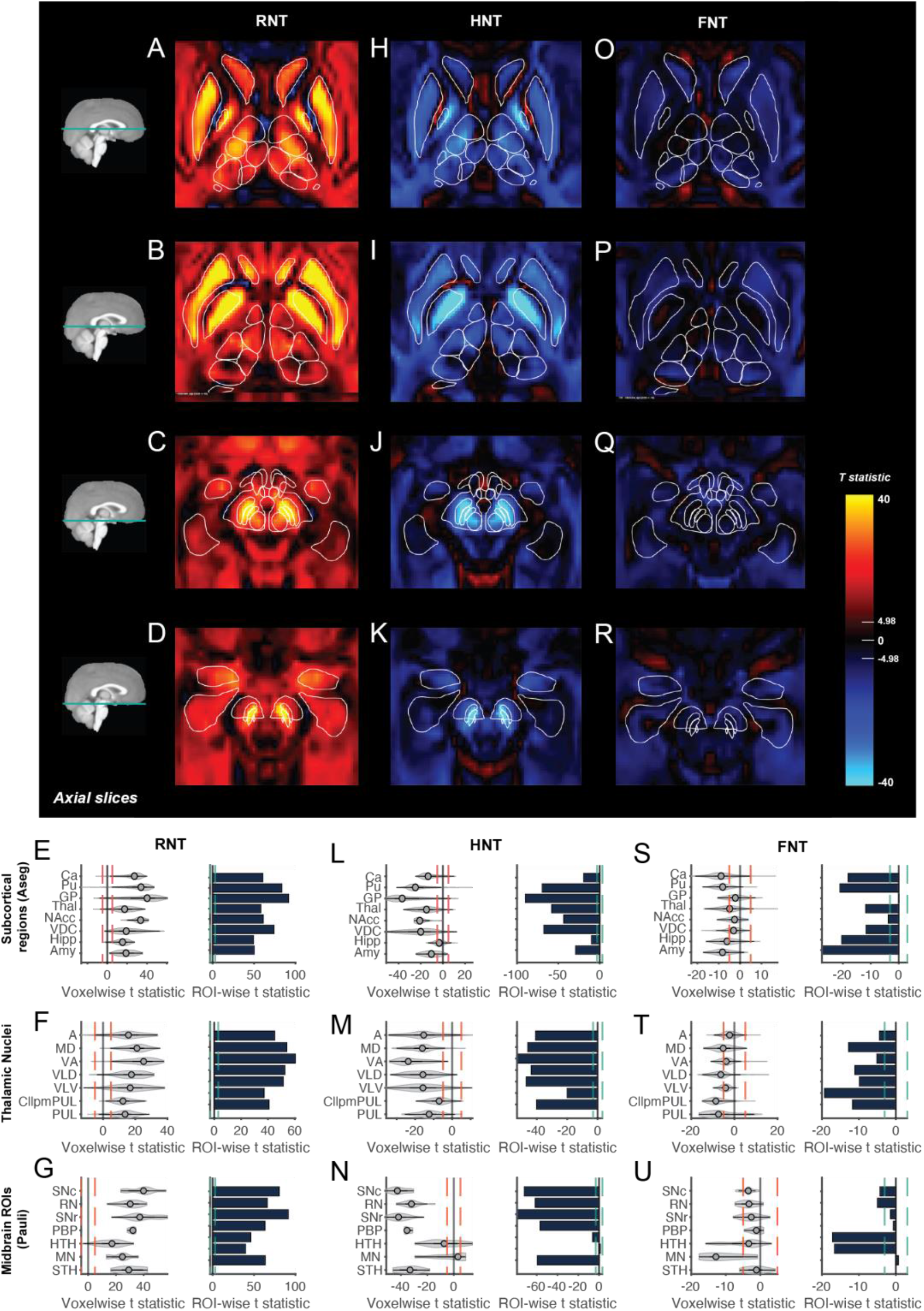
Age associations across diffusion compartments within subcortical regions. Voxelwise t-statistics for the association between age and RNT (A-D), HNT (H-K) and FNT (O-R) across different axial brain slices moving from superior (top) to inferior (bottom). Effects are unthresholded. Voxelwise Bonferroni corrected significance threshold (|t|=4.98) is marked on the colorbar. Outlines of the Aseg, Pauli and Najdenovska ROIs are overlaid. Violin plots show the distribution of voxelwise age associations in each ROI for each RSI metric. Red dotted lines show voxelwise Bonferroni corrected significance threshold. Bar plots show t-statistics from ROI analyses for the mean RSI metrics from each subcortical ROI. Green dotted line shows ROI Bonferroni corrected significance threshold (|t|=3.08). Plots are shown for RNT (E-G), HNT (L-N) and FNT (S-U). Subcortical ROI abbreviations are outlined in Supplementary Table 3.

Voxelwise age associations were extracted for each WM fiber tract in order to determine the distribution of effects within these regions. Violin plots highlight the heterogeneity of age-related changes in these RSI metrics within the WM fibers (Figure 4I-K). Moreover, these figures demonstrate the proportion of voxels above and below the conservative Bonferroni corrected threshold for whole-brain voxelwise statistical significance (red dotted line). ROI analyses were also conducted on the mean RSI metric within each fiber tract. For both RNI and RND, all WM fiber tracts showed significant positive age associations. The most significant ROI age effects for RNI were for the bilateral SCS, right SLF, and right CgH. The most significant ROI age effects for RND were for the bilateral SCS, and right ATR. For RDF, the most significant ROI age associations were found for the forceps major (FMaj), right inferior longitudinal fasciculus (ILF) and the right uncinate fasciculus (UF). Supplementary tables 8-10 show summary statistics for the voxelwise and ROIwise analyses within each WM fiber tract for RNI, RND and RDF.

Reflective of the RNT results, for age associations with RNI in particular, WM tracts with voxels near to or innervating subcortical regions appeared to show the greatest heterogeneity and largest voxelwise age associations, particularly the ATR, SCS and SIFC. This can clearly be seen for the SCS, where voxels in inferior portions of the tract overlapping with the putamen (Pu) ROI showed greater positive associations than voxels superior to the putamen within the SCS (supplementary figure 3A). Voxels that showed greater associations for RNI and RND within the SCS and overlaying with the Pu showed diffusion primarily in the anterior-posterior (green) direction, whereas more dorsal voxels showed diffusion primarily oriented in the dorsal-ventral (blue) direction as expected for diffusion along the SCS. This suggests greater age-related changes in restricted diffusion in voxels where the SCS potentially innervates the Pu. Similarly, along the ATR, age associations were greater in voxels overlapping with the thalamus and lower in voxels closer to the forceps minor (Fmin) (supplementary figure 3B). The difference in age associations between the FMaj and Fmin highlights a posterior-anterior gradient of development across the WM.

There were no significant voxelwise age-by-sex interaction effects for RNI, RND or RDF at the Bonferroni corrected significance threshold. Voxelwise age associations in a model without an age-by-sex interaction were highly correlated with a model including the interaction term (supplementary figure 2D-F). All main effects presented are from models without an age-by-sex interaction.

### Age associations in subcortical regions across the different RSI compartments

RNT was positively and highly significantly associated with age across subcortical regions, particularly within the basal ganglia. The only negative associations were found in voxels along the border between the Pu and globus pallidus (GP) and along the border of the caudate (Ca). The inverse relationship was found for HNT, as expected given the normalization of the RSI metrics. Voxelwise age effects were heterogeneous in magnitude across and within subcortical regions. Voxelwise FODs, averaged across participants, show the orientation structure of diffusion in each voxel and are colored based on the mean diffusion direction (Supplementary Figure 4). There was clear variability in the orientation structure of diffusion within gross subcortical ROIs and the surrounding WM, which likely contributes to the variability in voxelwise effects within these regions. We registered external subcortical atlases to our ABCD atlas in order to create finer subcortical parcellations to localize age-related effects within large subcortical structures. These included midbrain nuclei (Pauli et al., 2018) and thalamic nuclei (Najdenovska et al., 2018).

Voxels in the GP, Pu, the surrounding WM between and ventral to these structures, and voxels within the ventral diencephalon (VDC) showed the largest age associations with RNT and HNT (Figure 5A-N). Increases in RNT with age were found across the thalamic nuclei (Figure 5A,B,H,I) with the largest associations in more anterior nuclei. Voxelwise associations were the most heterogeneous within the VDC (Figure 5E,L). Within the VDC, voxels showing the largest RNT age associations were found within the substantia nigra pars compacta (SNpc), substantia nigra pars reticulata (SNpr) and the red nucelus (RN) (Figure 5C,D,J,K). ROI analyses, reflecting age associations with mean RSI metrics within each region, showed significant associations across all ROIs for RNT and HNT (except for the age association with HNT in the mamillary nucleus). There were limited age associations with FNT across subcortical regions. Supplementary tables 5-7 show summary statistics for the voxelwise and ROIwise analyses within each subcortical ROI for RNT, HNT and FNT.

**Figure 5.**
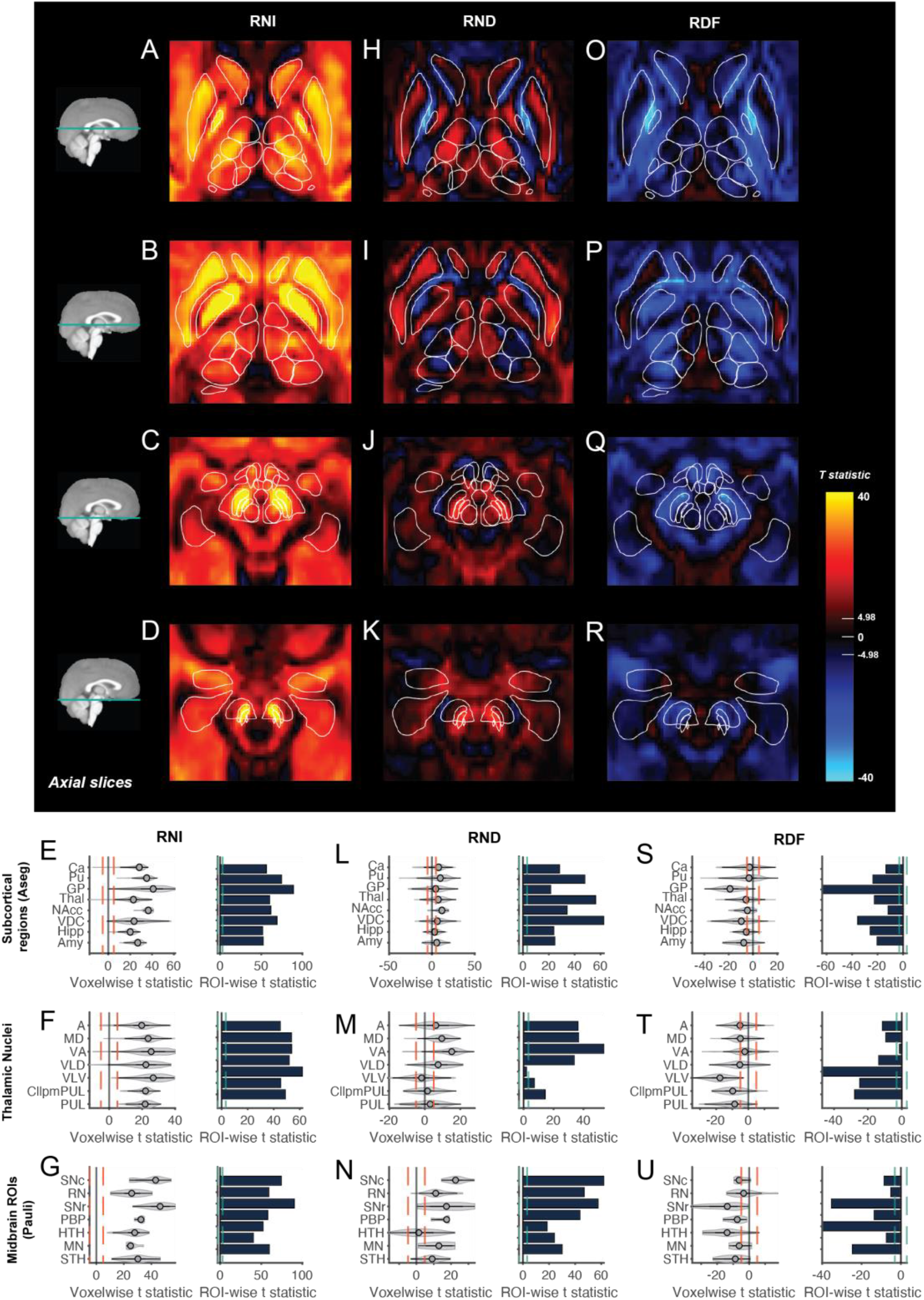
Age associations within the restricted compartment across subcortical regions. Voxelwise t-statistics for the association between age and RNI (A-D), RND (H-K) and RDF (O-R) across different axial brain slices moving from superior (top) to inferior (bottom). Effects are unthresholded. Voxelwise Bonferroni corrected significance threshold (|t|=4.98) is marked on the colorbar. Outlines of the Aseg, Pauli and Najdenovska ROIs are overlaid. Violin plots show the distribution of voxelwise age associations in each ROI for each RSI metric. Red dotted lines show voxelwise Bonferroni corrected significance threshold. Bar plots show t-statistics from ROI analyses for the mean RSI metrics from each subcortical ROI. Green dotted line shows ROI Bonferroni corrected significance threshold (|t|=3.08). Plots are shown for RNI (E-G), RND (L-N) and RDF (S-U). Subcortical ROI abbreviations are outlined in Supplementary Table 2.

### Age associations in subcortical regions within the restricted compartment

RNI was positively associated with age across all subcortical regions with the largest effects within the basal ganglia, particularly the GP, Pu and SN (Figure 6A-G). Voxelwise age associations within the thalamus (Thal) were largest in more anterior nuclei particularly along the lateral edge of the ventral anterior (VA) and ventro-lateral-ventral (VLV) nuclei (Figure 6A,B). ROI analyses showed a similar magnitude of effects across the thalamic nuclei for RNI. Age-related changes in RND were smaller in magnitude than RNI and more heterogeneous across voxels within and around subcortical regions (Figure 6H-N). The largest voxelwise age associations were primarily in voxels in the midbrain region, particularly the SN and RN, as well as the Pu, GP and Thal. Those regions also showed the largest ROI age associations. Negative age associations with RND were found in voxels along the border between the Pu and GP and along the border of the Ca (figure 6H,I); these effects were also seen for RNT. Within the Thal, there were a number of voxels within the anterior (A), VLV and pulvinar (P) nuclei that showed no significant age association. The largest positive age associations were found in the A, VA and medial dorsal (tMD) nuclei (Figure 6M). The heterogeneity in age effects across the Thal was particularly clear when looking at the ROI analysis for each nucleus. Across the Thal and VDC, the largest effects seemed to occur in voxels with diffusion occurring primarily within the anterior-posterior direction (Supplementary Figure 4). The most significant associations with RDF were in the GP, posterior nuclei of the thalamus, the substantia nigra pars reticulata, the hypothalmus and voxels along the border between the Pu and GP(Figure 6O-U). These were the regions that showed the largest difference in age associations between RNI and RND highlighting that the relative proportion of directional to isotropic diffusion within the restricted compartment decreased with age in these regions. Supplementary tables 8-10 show summary statistics for the voxelwise and ROIwise analyses within each subcortical ROI for RNI, RND and RDF.

Supplementary Figure 5 shows images of the most significant voxelwise age associations zoomed in on specific ROIs in order to highlight examples of how these associations occurred in voxels with particular diffusion orientation. Supplementary Figure 5A shows a coronal view of an area of RNI age associations extending ventral to the GP with diffusion occurring primarily in the lateral-medial (L-M) direction, which is likely to represent the anterior commissure; however, this location is difficult to distinguish from the ventral pallidum (VP), which sits below the anterior commissure. When looking in the sagittal view (supplementary figure 5B), we can see that these associations extend through the ventral striatum and head of the caudate. These effects appear to occur in voxels with diffusion in both the L-M and anterior-posterior (A-P) direction. The largest RNI and RND age effects in the VDC were seen in voxels with diffusion primarily in the A-P direction (Supplementary Figure 5C). Across the Thal, RNI and RND effects were larger in anterior nuclei where diffusion was also primarily in the A-P direction and along the intersection between the A, VA and VLV nuclei with diffusion in multiple directions (Supplementary Figure 5D,E).

### Age associations with DTI metrics

Supplementary figures 6-7 show the voxelwise and ROIwise age associations with MD and FA from the diffusion tensor model. Supplementary tables 11-12 show summary statistics for the voxelwise and ROIwise analyses within each WM fober tract and subcortical ROI for MD and FA. In general, there was a strong but inverse correspondence between the MD and RNI age associations across the WM and subcortical regions. However, there were subtle differences in the magnitude of effects across ROIs highlighting the different models used to estimate these measures. There were larger differences between the FA and RND associations. Namely, the magnitude of the FA associations was much smaller than RND, such that a larger sample would be required to detect age-related FA associations. However, the pattern of associations across the brain was similar.

## DISCUSSION

In this study, we have shown highly statistically significant age associated increases in restricted (primarily intracellular) diffusion across WM and subcortical GM in a large sample (n=8,086) of children from 9 to 14 years. This is the largest study to date measuring age associations in diffusion metrics at this age and utilizing novel RSI measures. Across both gray and WM, increasing age was associated with an increase in the proportion of restricted diffusion, RNT, and a decrease in the proportion of hindered, HNT, and free water, FNT. The largest age-related changes were found within the basal ganglia, namely the GP, and the VDC. Within the restricted compartment, the proportion of restricted isotropic diffusion, RNI, increased at a greater rate with age than directional diffusion, RND, resulting in a relative increase in the isotropic compared to directional signal fraction across the brain. These differences were most pronounced in the GP, posterior nuclei of the Thal and the midbrain nuclei. Voxelwise age associations were highly variable within subcortical regions and WM fiber tracts. Within subcortical regions in particular, the pattern of age associations appeared to follow changes in the orientation of the diffusion. This suggests that we can identify distinct age associations within subcomponents of subcortical structures that may be associated with differing functional circuits as indicated by differences in cytoarchitecture. This highlights the benefit of measuring voxelwise compared to ROI-wise associations and utilizing high resolution parcellations of subcortical structures that reflect known histologic and functional subdivisions within deep gray matter nuclei such as the Thal.

### White Matter associations with age

There was a robust increase in the proportion of restricted to hindered and free water from 9-14 years across the WM, which was associated with an increase in both isotropic and directional diffusion. In general, RNI showed more widespread effects across the WM compared to RND, which showed larger age associations along the centers of WM tracts where axonal coherence is highest. In many voxels, increases in both RNI and RND reflected increases in the overall proportion of the restricted signal fraction relative to the hindered and free water compartments. To tease apart differences in the magnitude of the RNI and RND associations with age we estimated the relative proportion of restricted directional diffusion over the total restricted signal fraction, RDF. The negative voxelwise associations between RDF and age highlighted that the proportion of restricted directional diffusion within the restricted compartment was decreasing with age i.e. directional diffusion was increasing with age at a lower rate than isotropic diffusion. This was seen in both GM and WM. There were some regions, such as the Pu, that showed limited RDF effects, highlighting that there were similar increases in the restricted isotropic and directional fractions in this region. Increases in isotropic diffusion can be driven by both an increase in the size or number of structures with a spherical or compact shape, and multiple cylindrical structures in the same voxel oriented such that anisotropic diffusion is occurring in all directions appearing as isotropic. Therefore, an increase in the complexity of neuronal connections and crossing fibers at angles smaller than can be resolved could lead to an increase in the relative contribution of isotropic compared to directional diffusion within the restricted compartment with age. This is in contrast to previous work using the NODDI model in which orientation dispersion (a measure of the degree of dispersion of neurites) was not found to increase with age (Chang et al., 2015; Genc, Seal, et al., 2017; Mah et al., 2017). This may reflect increased statistical power in this study to detect an association or key differences in the diffusion model and metrics estimated. Future work comparing multiple multi-compartment diffusion models using large developmental samples will be required to tease apart these differences.

The largest and most heterogeneous voxelwise effects across WM fibers, particularly for RNI, were within the ATR, SCS and SIFC, such that significant ROI associations appeared to be driven by voxels within or near subcortical structures. For the SCS, the tractography used to generate the SCS tract ROI included termination points in the striatum (Hagler et al., 2019), therefore the overlap of the SCS ROI and the Pu ROI is likely indicating voxels in which the SCS is innervating the Pu. The greater age associations in this region suggests there may be greater age-related changes in WM microstructure at this innervation point. The same could be seen for the ATR; in the original tractography all streamlines for the ATR were set to terminate on one end of the thalamus and not pass through the thalamus. The greater age-related associations in anterior thalamic nuclei innervated by the ATR may highlight specific refinement of particular circuits involving these nuclei. These analyses highlight the importance of measuring voxelwise associations to avoid the misleading impression of homogeneity of effects across the entirety of WM tracts. This has been eloquently shown previously using a similar model of intracellular diffusion (Lynch et al., 2020).

In the corpus callosum (CC), RNI and RND both increased with age, however, the magnitude of the age associations differed along the posterior-anterior axis. Voxels in the Fmaj, showed a greater age effect than voxels in the Fmin, which connects the lateral and medial surfaces of the frontal lobes and is the frontal portion of the CC. Our results support previous evidence from other developmental samples showing a greater age effect of intracellular diffusion metrics in the splenium or forceps major compared to the genu or forceps minor (Geeraert et al., 2019; Genc, Seal, et al., 2017; Mah et al., 2017) and the more extended development of frontal-temporal connections (Genc, Seal, et al., 2017; C. Lebel et al., 2008; Catherine Lebel & Beaulieu, 2011; Tamnes et al., 2010). This mirrors the posterior-anterior sequence of myelination in developing infants (Bird et al., 1989; Kinney et al., 1988), suggesting differences in the time-course of myelination across the CC may be contributing to the effects here. In addition, Genc et al found that from 4 to 19 years, age showed a greater positive association with apparent fiber density (a measure of the intracellular volume fraction) in posterior relative to anterior portions of the CC (Genc et al., 2018). This suggests that changes in axonal diameter and/or myelination, that can contribute to increases in the restricted volume fraction, likely occur at a different rate depending on the location in the CC. Different sections of the CC connect different cortical regions within distinct functional networks. Nonuniformity in the development of these interhemispheric connections may reflect age-dependent maturation of cognitive and behavioral processes. More protracted developmental changes in frontal circuitry may underpin the later development of cognitive control in adolescence and become more prominent as the children get older (Casey et al., 2008).

### Subcortical associations with age

Previous studies have highlighted significant changes in FA and MD across subcortical regions (Baron Nelson et al., 2019; Lebel et al., 2008; Simmonds et al., 2014), with regions of the basal ganglia showing greater percentage change from 5-30 years than many WM tracts (Lebel et al., 2008), in agreement with the results reported here. From 8-13 years, Mah et al (2017) found that NDI from the NODDI model, a measure of the intracellular volume fraction, showed the largest percent increase in the GP (10-13% change) followed by the Pu, hippocampus, amygdala and Thal (3-7% change) and found no age association in the Ca. Although the RSI and NODDI models are very different, NDI, similar to RNT, captures the total intracellular volume fraction in a voxel. As the intracellular volume fraction increases in a voxel, the magnitude of water displacement reduces, thereby decreasing MD. Indeed, NDI has previously been shown to correlate negatively with MD (Zhang et al., 2012), and in the current study MD showed age associations in the opposite direction to RNT (as expected). Indeed, our RNT results were very similar to the NDI effects reported by Mah et al., apart from a significant age association in the caudate. This may reflect the greater sensitivity of the RSI model parameters to age-related changes in cytoarchitecture of the caudate and/or increased statistical power in this study to detect an association.

By using voxelwise analyses we were able to measure the heterogeneity of developmental effects within subcortical regions highlighting the benefit of using voxelwise compared to ROI-wise analyses. There was a clear pattern of age associations across the different thalamic nuclei, particularly for RND. Najdenovska et al (2018) generated the thalamic nuclei ROIs by clustering contiguous voxels with similar orientation microstructure (determined by the FODs) validating their results against a histological atlas (Najdenovska et al., 2018). When overlaying these ROIs on the average FODs measured in our sample (Supplementary Figure 4), we could see that the boundaries of the different nuclei indeed adhered to changes in the primary orientation of diffusion. Within anterior nuclei (A, VA, tMD), where age associations were greatest for RNI and RND, diffusion primarily occurred in the anterior-posterior orientation (green), whereas within posterior nuclei (VLV, VLD, C, P), diffusion primarily occurred within the lateral-medial (red) orientation. The tMD nucleus of the thalamus is reciprocally interconnected with the prefrontal cortex and receives input from striatal, medial temporal, midbrain and basal forebrain structures (Groenewegen, 1988; Groenewegen et al., 1993; Ray & Price, 1993; Tanaka, 1976; Tobias, 1975; Vertes et al., 2015). It is well positioned to play a modulatory role within fronto-striatal-thalamo-cortical circuits thought to be important for several cognitive and emotional processes (Haber & Calzavara, 2009; Mitchell & Chakraborty, 2013; Ouhaz et al., 2018). Structural and functional connectivity of these thalamo-cortical connections has been shown to increase across childhood and adolescence (Alkonyi et al., 2011; Fair, 2010), and is thought to underpin behavioral changes in cognitive control and emotional reactivity during adolescence.

Moreover, there were highly significant age associations with RNI in the region ventral to the GP and Ca, which encompasses both the ventral pallidum, ventral striatum (nucleus accumbens and olfactory tubercle) and bed of the nucleus stria terminalis (often referred to as the extended amygdala, as well as the anterior commissure (Zaborszky et al., 2015). These regions are highly interconnected with subcortical and cortical regions, particularly in frontal cortex, creating circuits integral for incentive-based learning, reward processing and decision-making (Barkley-Levenson & Galván, 2014; Delgado, 2007; Haber & Knutson, 2010). Microstructural changes within the thalamus and the ventral forebrain may be indicative of the refinement of these circuits in late childhood.

There were also statistically robust and heterogeneous associations within the VDC. The VDC is a group of structures that are poorly defined on T1w imaging, however, by calculating the mean voxelwise FODs across subjects, we could clearly see variability in the orientation of diffusion within this large region highlighting the presence of potentially distinct cytoarchitecture. Changes in the orientation of the FODs also appeared to adhere to estimated outlines of finer subcortical parcellations that include many of the nuclei within the VDC from the Pauli atlas (Pauli et al., 2018). Indeed, the strongest and most significant associations between age and RNI and RND were in voxels oriented primarily in the anterior-posterior direction within and around the SN adjacent to the ventral tegmental area (VTA), which may reflect microstructural changes within the extensive dopaminergic projections from these regions to the basal ganglia and medial forebrain. Fibers from the SN and striatum also directly innervate the lateral edge of the VA nucleus where we see high isotropy and crossing diffusion orientations (Kultas-ilinsky & Ilinsky, 1990; Sakai et al., 1998). Our findings may signal increased innervation of the Thal from the SN and/or striatum and/or increased myelination of axons in these pathways. This further suggests that these findings may reflect maturation of cortico-striatal-thalamic pathways that may be important for processes of motor control, cognition and self-regulation that are developing in this age range. Indeed, FA in the internal capsule, basal ganglia and Thal has been shown to partially mediate improved performance on particular cognitive tasks with increasing age from 9-12 years (Baron Nelson et al., 2019).

Voxels within the GP showed the largest age associations with RNI, followed by the Pu. These basal ganglia regions form part of parallel frontal, temporal and parietal cortical circuits that are involved in a number of cognitive and motor functions (Alexander et al., 1986; Middleton & Strick, 2000). Overall mean restricted isotropic diffusion across subjects was much larger in the GP than other subcortical regions, which may reflect higher myelin content in the GP compared to the Pu (Lanciego et al., 2012). Moreover, throughout adolescence, there is a large increase in iron concentration, estimated by increased susceptibility on T2* weighted imaging, within the GP (Larson et al, 2020). This iron related effect has been shown to correlate with DTI metrics in these deep GM structures (Pfefferbaum et al., 2010); therefore, increasing iron accumulation may be contributing to the RNI developmental effects that we observe in this region. Further research is required to determine the extent to which iron content effects the estimation of diffusion metrics.

### Microstructural changes underlying alterations in restricted diffusion

There are several biological processes that may increase restricted diffusion in the WM, such as increasing myelination, neurite density and/or axon coherence. Increasing myelination reduces the permeability of myelinated axons and decreases the volume of the extracellular space in a voxel increasing the restricted signal fraction. Previous studies using NODDI have shown age-related changes in NDI (a measure of the intracellular volume fraction similar to RNT) in a similar age range in the WM (Chang et al., 2015; Genc, Seal, et al., 2017; Mah et al., 2017) and increases in NDI have been associated with both increases in the myelin volume fraction using MRI (Geeraert et al., 2019) and post-mortem histology from patients with demyelination (Grussu et al., 2017). In general, there has been a lack of evidence for increasing myelination in late childhood as measured with magnetization transfer (Geeraert et al., 2019; Moura et al., 2016; Pangelinan et al., 2016). However, given histological findings that myelination continues into adulthood (Benes, 1989; Yakovlev & Lecours, 1967), these myelin changes may be very subtle, and require large sample sizes and/or longitudinal studies, such as this, to detect. Little is known about how changes in myelination directly impact RSI measures specifically; however, in a demyelinated genetic mouse model, regions with reduced myelin showed reduced intraneurite volume fraction, estimated using a similar spherical deconvolution method (Kaden et al., 2016). Increased myelination with development may underlie the age effects observed here; however, this is only one of many biological processes that can alter restricted diffusion, as described in Table 1, therefore further work is needed to elucidate the exact neurobiological processes contributing to the effects reported here.

Subcortical gray matter has a more complex cytoarchitecture than cortical gray matter and WM fiber bundles consisting of many cell bodies of stellate shape, dendrites, terminal arbors and synapses that do not follow a coherent structure. Nevertheless, examination of the mean estimated FODs from the RSI model showed complex orientation structure within these regions. Developmental increases in the restricted signal fraction within the deep gray matter nuclei could be driven by increases in myelination, neurite density, dendritic sprouting or increases in cell size less than the typical diffusion length scale. These microstructural possibilities are outlined in Table 1. As mentioned previously, increases in restricted isotropic diffusion measured by RSI can be modulated by both structures with a spherical or compact shape less than the diffusion length scale, such as neuronal cell bodies or microglia, within which diffusion is isotropic, and also by multiple cylindrical structures in the same voxel oriented such that anisotropic diffusion is occurring in all directions. By using RSI we can detect differences in diffusion orientation at a much finer resolution than with the tensor model allowing us to more precisely understand the underlying microstructural changes occurring during late childhood and adolescent development.

### DTI vs RSI

In the current study, age-related changes in RNT were very similar (but opposite in sign) to those observed in MD, estimated from the diffusion tensor model. As diffusion becomes more restricted, MD within a voxel will decrease. Our analyses showed greater age-related changes in MD compared to FA from 9-14 years, whereas previous studies have shown relatively greater percent changes in FA compared to MD across WM tracts in particular (Krogsrud et al., 2016b; C. Lebel et al., 2008). These studies are difficult to compare directly due to different sample sizes and age ranges. Furthermore, most studies estimate MD using diffusion MRI data acquired at lower b-values (below b=2000 s/mm^2^) than the current study (which includes many directions at b=3000 s/mm^2^). At higher b-values, around b≥3000s/mm^2^, the signal from the hindered compartment is attenuated and the measured diffusion signal is dominated by the restricted compartment. If maturational changes are predominantly occurring within the restricted compartment our DTI measures may be more sensitive to age-related changes than DTI metrics from studies with lower b-value acquisitions. Further work empirically comparing age-related changes on DTI metrics calculated with different acquisition parameters is required.

### Limitations

Although the effects in this study are highly significant due to the large sample size, the magnitude of the voxelwise effects is very small. This has been observed across large-scale imaging studies making it clear that published effects are inflated by small sample sizes and publication bias (Dick et al., 2021). Using large sample sizes we are now able to estimate effect sizes with much greater precision. Small effects are perhaps not surprising given that causal associations among variables are highly complex and a single association is unlikely to be very large. These large sample studies can provide new norms for expected effect sizes. Moreover, the data in the current study only included two timepoints, therefore we were unable to disentangle regional differences in the developmental trajectory of these RSI metrics from 9-14 years. Imaging data were also rank normalized to make the assumptions of normality in the statistical analysis valid, therefore we did not measure non-linear age associations as the non-linear aspect of the rank normalization could introduce apparent non-linearities. With future ABCD Study data releases with more longitudinal time-points, future work should aim to map the shape of developmental trajectories of these RSI metrics. Finally, given the increased T2 shortening of the diffusion signal with increased iron concentrations, future work should aim to understand the implications of this for diffusion models.

### Conclusions

This is the largest study to date measuring longitudinal age associations with diffusion metrics across the brain. We have demonstrated highly significant increases in the proportion of restricted diffusion across the WM and within deep gray matter nuclei. The heterogeneity of effects along WM tracts and within subcortical GM structures highlights the importance of voxelwise analyses to provide a more fine-grained understanding of how the brain is changing with age. Given the importance of both subcortical-subcortical and subcortical-cortical circuitry in the development of multiple cognitive and behavioral dimensions during this period (Casey et al., 2016), robust microstructural changes occurring in subcortical regions and associated WM tracts may indicate important refinement of these developing circuits. Understanding whether individual differences in the age-related structural covariance of these measures associates with differential behavioral profiles will provide a promising new avenue for future research.

## ACKNOWLEDGEMENTS

The authors wish to thank the youth and families participating in the Adolescent Brain Cognitive Development (ABCD) Study and all ABCD staff involved in data collection and curation. Dr. Dale reports that he was a Founder of and holds equity in CorTechs Labs, Inc., and serves on its Scientific Advisory Board. He is a member of the Scientific Advisory Board of Human Longevity, Inc. He receives funding through research grants from GE Healthcare to UCSD. The terms of these arrangements have been reviewed by and approved by UCSD in accordance with its conflict of interest policies.

## Footnotes

1 The free normalized total signal fraction (FNT) is equivalent to the free normalized isotropic signal fraction (FNI) from Release 4.0 of the ABCD Study. These are equivalent because there is no directional component to the free water compartment, but was renamed here for consistency with RNT and HNT.

