## Supplementary Material for "Microstructural development from 9-14 years: evidence from the ABCD Study"

### SUPPLEMENTARY TABLES & FIGURES

| | | |
|---|---|---|
| **Supplementary Table 1** | Quality control (QC) metrics for the imaging data | Pg 2 |
| **Supplementary Table 2** | Fiber tracts automatically labeled by AtlasTrack | Pg 3 |
| **Supplementary Table 3** | Subcortical ROIs labelled using 3 different methods | Pg 4 |
| **Supplementary Table 4** | Estimated mean voxelwise scanner and software version effects | Pg 5 |
| **Supplementary Table 5** | Summary statistics for the RNT age associations | Pg 6 |
| **Supplementary Table 6** | Summary statistics for the HNT age associations | Pg 7 |
| **Supplementary Table 7** | Summary statistics for the FNT age associations | Pg 8 |
| **Supplementary Table 8** | Summary statistics for the RNI age associations | Pg 9 |
| **Supplementary Table 9** | Summary statistics for the RND age associations | Pg 10 |
| **Supplementary Table 10** | Summary statistics for the RDF age associations | Pg 11 |
| **Supplementary Table 11** | Summary statistics for the MD age associations | Pg 12 |
| **Supplementary Table 12** | Summary statistics for the FA age associations | Pg 13 |
| **Supplementary Figure 1** | Voxelwise color-coded FA maps with labelled WM fiber tracts | Pg 14 |
| **Supplementary Figure 2** | Association between voxelwise age associations with and without including an age-by-sex interaction | Pg 15 |
| **Supplementary Figure 3** | Associations between age, RNI and RND in specific tracts | Pg 16 |
| **Supplementary Figure 4** | Voxelwise FODs averaged over participants within subcortical regions | Pg 17 |
| **Supplementary Figure 5** | Zoomed in images of the voxelwise age associations with RNI, RND and RDF and mean FODs across different brain slices | Pg 18 |
| **Supplementary Figure 6** | Associations between age and DTI metrics across the brain | Pg 19 |
| **Supplementary Figure 7** | Associations between age and DTI metrics within subcortical regions | Pg 20 |

**Supplementary Table 1.** Quality control (QC) metrics for the imaging data taken from the ABCD Study Release notes. All imaging data included in this study passed the dMRI include flag, the T1w include flag and had an incidental finding score less than 3 (incl_dmri_include==1 & imgincl_t1w_include==1 & mrif_score<3). The tables below highlight all the QC metrics that each scan had to fulfill to be included.

| T1w Criteria | Instrument | Element value |
|---|---|---|
| **T1 series passed rawQC** | mriqcrp103 | iqc_t1_ok_ser > 0 |
| **FreeSurfer QC not failed** | abcd_fsurfqc01 | fsqc_qc ~= 0 |
| **Derived results exist** | abcd_smrip202 | smri_t1w_scs_cbwmatterlh ~= NA |

| dMRI Criteria | Instrument | Element value |
|---|---|---|
| **dMRI series passed rawQC** | mriqcrp103 | iqc_dmri_ok_ser > 0 |
| **dMRI total number of repetitions** | mriqcrp103 | iqc_dmri_ok_nreps >= 103 *OR* (mri_info_manufacturer = Philips AND iqc_dmri_ok_ser >= 2 AND iqc_dmri_ok_nreps = 51) |
| **T1 series passed rawQC** | mriqcrp103 | iqc_t1_ok_ser > 0 |
| **dMRI B0 unwarp available** | abcd_auto_postqc01 | apqc_dmri_bounwarp_flag == 1 |
| **FreeSurfer QC not failed** | abcd_fsurfqc01 | fsqc_qc ~= 0 |
| **dMRI manual post-processing QC not failed** | abcd_dmriqc01 | dmri_postqc_qc ~= 0 |
| **dMRI registration to T1w** | abcd_auto_postqc01 | apqc_dmri_regt1_rigid < 17 |
| **dMRI dorsal cutoff score** | abcd_auto_postqc01 | apqc_dmri_fov_cutoff_dorsal < 47 |
| **dMRI ventral cutoff score** | abcd_auto_postqc01 | apqc_dmri_fov_cutoff_ventral < 54 |
| **Derived results exist** | abcd_drsip201 | dmri_rsirnd_fib_allfib ~= NA |

**Supplementary Table 2. Fiber tracts automatically labeled by AtlasTrack.** All tracts are separately labeled for left and right hemispheres except for Fmaj, Fmin, and CC (Hagler, et al., 2009). The Fmaj and Fmin are the occipital and frontal subsets of the CC respectively.

| **Abbrev** | **Fiber Name** | **Connected Brain Regions** |
|---|---|---|
| Fx | fornix | hippocampus & mammilary nuclei of hypothalamus |
| cCing | cingulate cingulum | cingulate gyrus & entorhinal cortex (cingulate portion) |
| pCing | parahippocampal cingulum | cingulate gyrus & entorhinal cortex (parahippocampal portion) |
| CST | corticospinal tract (pyramidal tract) | motor cortex & spinal cord |
| ATR | anterior thalamic radiations | thalamus & frontal lobe |
| UF | uncinate | inferior frontal lobe & anterior temporal lobe |
| ILF | inferior longitudinal fasciculus | occipital lobe & temporal lobe |
| IFOF | inferior frontal occipital fasciculus | occipital lobe & frontal lobe |
| Fmaj | forceps major | left occipital cortex & right occipital cortex |
| Fmin | forceps minor | left prefrontal cortex & right prefrontal cortex |
| CC | corpus callosum | left cortex & right cortex |
| SLF | superior longitudinal fasciculus | temporal and parietal lobes & frontal lobe |
| SCS | superior corticostriate | superior cortex & striatum |
| SIF | striatal inferior frontal cortex tract | inferior frontal cortex & striatum |
| IFSF | inferior frontal to superior frontal cortical tract | inferior frontal cortex & superior frontal cortex |

**Supplementary Table 3. Subcortical regions of interest (ROIs) labelled using 3 different methods.** Column 1) automatic segmentation using FreeSurfer 5.3 applied to each subject's T1 image in atlas space (Fischl et al., 2002); Column 2) registration of the Pauli atlas of subcortical nuclei to the multispectral atlas (Pauli et al., 2018); Column 3) registration of the the Najdenovska thalamic nuclei atlas to our data (Najdenovska et al., 2018).

| **FreeSurfer 5.3 segmentation**<br>**T1** | **Pauli, 2018**<br>**HCP T1 & T2** | **Najdenovska, 2018**<br>**HCP FODs** |
|---|---|---|
| Amygdala (Amg) | Extended amygdala (EA) | Anterior (A) |
| Hippocampus (Hipp) | Substantia nigra pars compacta (SNpc) | Ventral anterior (VA) |
| Putamen (Pu) | Substantia nigra pars reticulata (SNpr) | Mediodorsal (tMD) |
| Caudate (Ca) | Red nucleus (RN) | Ventral-latero-ventral (VLV) |
| Globus pallidus (GP) | Parabrachial pigmented nucleus (PBP) | Ventral-latero-dorsal (VLD) |
| Accumbens area (NAcc) | Hypothalamus (Hyp) | Central-latero-lateral-posterior-medial-pulvinar (C) |
| Thalamus (Thal) | Mamillary nucleus (MN) | Pulvinar (P) |
| Ventral Diencephalon (VDC) | Subthalamic nucleus (STN) | |

**Supplementary Table 4.** Estimated mean voxelwise scanner and software version effects for each imaging modality. Pseudo-$\Delta R^2$ estimates were calculated by taking the difference between the mean voxelwise pseudo-$R^2$ for the full model including all predictors and the mean voxelwise pseudo-$R^2$ for a reduced model without either the dummy coded scanner predictors or the dummy coded software version predictors.

| Modality | Scanner effect Pseudo-$\Delta R^2$ | Software version Pseudo-$\Delta R^2$ |
|---|---|---|
| **RNT** | 0.0175 | 0.0041 |
| **HNT** | 0.0140 | 0.0034 |
| **FNI** | 0.0194 | 0.0053 |
| **RNI** | 0.0298 | 0.0053 |
| **RND** | 0.0107 | 0.0031 |
| **RDF** | 0.0155 | 0.0040 |
| **FA** | 0.0141 | 0.0066 |
| **MD** | 0.0247 | 0.0069 |

**Supplementary Table 5. Summary statistics for the RNT age associations.** Statistics include the range of beta coefficients and t statistics within each ROI for the voxelwise analyses and the estimated beta coefficient, SE and t statistic the ROI analyses.

| | | age voxelwise effects | | | | age ROIwise effects | | |
|---|---|---|---|---|---|---|---|---|
| **Atlas** | **ROI** | $min\beta$ | $max\beta$ | $min\ t$ | $max\ t$ | $\beta$ | $SE$ | $t$ |
| **AtlasTrack** | CC | -0.0031 | 0.0057 | -15.64 | 37.81 | 0.011 | 0.00027 | 42.08 |
| **AtlasTrack** | Fmaj | -0.003 | 0.0038 | -15.64 | 23.71 | 0.011 | 0.00025 | 45.24 |
| **AtlasTrack** | Fmin | -0.0026 | 0.0033 | -11.86 | 16.29 | 0.005 | 0.00028 | 17.64 |
| **AtlasTrack** | R Fx | -0.0018 | 0.0045 | -8.68 | 20.55 | 0.0087 | 0.00035 | 24.79 |
| **AtlasTrack** | L Fx | -0.0016 | 0.0047 | -8.38 | 21.46 | 0.0065 | 0.00029 | 22.43 |
| **AtlasTrack** | R CgC | -0.0006 | 0.0051 | -3.79 | 28.87 | 0.011 | 0.00032 | 35.79 |
| **AtlasTrack** | L CgC | -0.0013 | 0.0057 | -6.59 | 30.13 | 0.011 | 0.00035 | 32.1 |
| **AtlasTrack** | R CgH | 0.0002 | 0.0048 | 0.98 | 23.87 | 0.016 | 0.00032 | 49.97 |
| **AtlasTrack** | L CgH | -0.00049 | 0.0049 | -2.42 | 25.35 | 0.016 | 0.00032 | 49.36 |
| **AtlasTrack** | R CST | -0.00083 | 0.0063 | -3.79 | 29.14 | 0.017 | 0.0003 | 55.93 |
| **AtlasTrack** | L CST | -0.0011 | 0.008 | -5.47 | 37.81 | 0.016 | 0.0003 | 55.29 |
| **AtlasTrack** | R ATR | -0.0021 | 0.01 | -9.94 | 46.82 | 0.017 | 0.00034 | 49.57 |
| **AtlasTrack** | L ATR | -0.0043 | 0.011 | -21.01 | 49.43 | 0.016 | 0.00035 | 44.6 |
| **AtlasTrack** | R Unc | -0.0017 | 0.0038 | -7.95 | 21.22 | 0.01 | 0.00025 | 40.6 |
| **AtlasTrack** | L Unc | -0.00062 | 0.0036 | -3.17 | 25.35 | 0.012 | 0.00025 | 46.69 |
| **AtlasTrack** | R ILF | -0.00014 | 0.004 | -0.83 | 23.22 | 0.013 | 0.00031 | 41.3 |
| **AtlasTrack** | L ILF | 4.90E-06 | 0.0041 | 0.03 | 23.15 | 0.012 | 0.00028 | 45.35 |
| **AtlasTrack** | R IFO | -0.0017 | 0.0047 | -7.95 | 22.22 | 0.013 | 0.00026 | 48.43 |
| **AtlasTrack** | L IFO | -0.0018 | 0.0075 | -7.68 | 33.42 | 0.013 | 0.00026 | 50.72 |
| **AtlasTrack** | R SLF | 0.00067 | 0.005 | 3.4 | 28.07 | 0.016 | 0.00026 | 63.55 |
| **AtlasTrack** | L SLF | 0.0012 | 0.0048 | 6.26 | 26.37 | 0.015 | 0.00027 | 55.88 |
| **AtlasTrack** | R SCS | 0.00019 | 0.0098 | 0.98 | 45.73 | 0.024 | 0.00032 | 74.14 |
| **AtlasTrack** | L SCS | -0.00038 | 0.01 | -1.75 | 45.82 | 0.024 | 0.00032 | 74.11 |
| **AtlasTrack** | R SIFC | -0.0012 | 0.01 | -5.69 | 45.57 | 0.014 | 0.00032 | 44.46 |
| **AtlasTrack** | L SIFC | -0.0018 | 0.0093 | -7.68 | 43.17 | 0.017 | 0.00029 | 57.05 |
| **AtlasTrack** | R IFSFC | 0.00032 | 0.0058 | 1.69 | 29.14 | 0.017 | 0.00028 | 60.95 |
| **AtlasTrack** | L IFSFC | 0.00021 | 0.0051 | 1.21 | 26.93 | 0.015 | 0.00028 | 54.51 |
| **Freesurfer Aseg** | Ca | -0.0021 | 0.0089 | -11.23 | 39.94 | 0.019 | 0.00031 | 60.77 |
| **Freesurfer Aseg** | Pu | -0.0059 | 0.01 | -26.19 | 47.28 | 0.027 | 0.00032 | 83.58 |
| **Freesurfer Aseg** | GP | -0.0032 | 0.012 | -13.5 | 60.65 | 0.03 | 0.00033 | 91.75 |
| **Freesurfer Aseg** | Thal | -0.0029 | 0.0084 | -13.91 | 38.21 | 0.022 | 0.00037 | 58.25 |
| **Freesurfer Aseg** | NAcc | 0.003 | 0.009 | 13.96 | 41.64 | 0.022 | 0.00036 | 61.13 |
| **Freesurfer Aseg** | VDC | -0.00055 | 0.013 | -2.67 | 57.18 | 0.028 | 0.00037 | 74.12 |
| **Freesurfer Aseg** | Hipp | -0.00075 | 0.0063 | -3.75 | 27.55 | 0.017 | 0.00034 | 49.43 |
| **Freesurfer Aseg** | Amy | 0.00046 | 0.0082 | 2.17 | 35.63 | 0.022 | 0.00043 | 50.21 |
| **Najdenovska** | A | -0.0029 | 0.0076 | -13.54 | 33.81 | 0.018 | 0.0004 | 44.98 |
| **Najdenovska** | MD | -0.00045 | 0.0081 | -1.87 | 35.4 | 0.02 | 0.00038 | 54.02 |
| **Najdenovska** | VA | -0.0016 | 0.0082 | -7.17 | 37.85 | 0.023 | 0.00038 | 60.45 |
| **Najdenovska** | VLD | -0.0017 | 0.0071 | -8.49 | 31.64 | 0.017 | 0.00032 | 52.63 |
| **Najdenovska** | VLV | -0.0028 | 0.0084 | -12.07 | 38.21 | 0.021 | 0.00041 | 51.5 |
| **Najdenovska** | CllpmPUL | -0.00048 | 0.0059 | -2.23 | 26.87 | 0.011 | 0.00031 | 37.11 |
| **Najdenovska** | PUL | -0.0021 | 0.0057 | -9.57 | 28.74 | 0.014 | 0.00034 | 40.82 |
| **Pauli** | SNc | 0.0052 | 0.012 | 23.52 | 56.09 | 0.031 | 0.00039 | 80.48 |
| **Pauli** | RN | 0.003 | 0.0096 | 13.66 | 42.32 | 0.027 | 0.00041 | 66.05 |
| **Pauli** | SNr | 0.0037 | 0.013 | 16.92 | 57.18 | 0.033 | 0.00036 | 91.36 |
| **Pauli** | PBP | 0.0061 | 0.0079 | 28.13 | 34.43 | 0.026 | 0.00042 | 63.26 |
| **Pauli** | HTH | -0.0008 | 0.0073 | -3.67 | 32.8 | 0.018 | 0.0004 | 46.06 |
| **Pauli** | MN | 0.0027 | 0.0081 | 13.13 | 36.12 | 0.018 | 0.00045 | 39.72 |
| **Pauli** | STH | 0.0036 | 0.0099 | 16.32 | 42.76 | 0.027 | 0.00043 | 63.69 |

**Supplementary Table 6. Summary statistics for the HNT age associations.** Statistics include the range of beta coefficients and t statistics within each ROI for the voxelwise analyses and the estimated beta coefficient, SE and t statistic the ROI analyses.

| | | age voxelwise effects | | | | age ROIwise effects | | |
|---|---|---|---|---|---|---|---|---|
| **Atlas** | **ROI** | $min\beta$ | $max\beta$ | $min\ t$ | $max\ t$ | $\beta$ | $SE$ | $t$ |
| **AtlasTrack** | CC | -0.0063 | 0.0032 | -37.16 | 13.18 | -0.008 | 0.00032 | -25.05 |
| **AtlasTrack** | Fmaj | -0.0042 | 0.0017 | -24.34 | 8.04 | -0.0093 | 0.00029 | -31.64 |
| **AtlasTrack** | Fmin | -0.0033 | 0.0029 | -16.99 | 13.01 | -0.003 | 0.00031 | -9.7 |
| **AtlasTrack** | R Fx | -0.0052 | 0.0012 | -26.64 | 6.18 | -0.011 | 0.00044 | -24.89 |
| **AtlasTrack** | L Fx | -0.0043 | 0.0011 | -20.64 | 4.46 | -0.011 | 0.00042 | -27.14 |
| **AtlasTrack** | R CgC | -0.0055 | 0.00054 | -28.84 | 2.66 | -0.01 | 0.00034 | -29.9 |
| **AtlasTrack** | L CgC | -0.006 | 0.0024 | -29.06 | 10.68 | -0.0092 | 0.00038 | -24.46 |
| **AtlasTrack** | R CgH | -0.0054 | -0.00057 | -26.25 | -2.87 | -0.014 | 0.00036 | -39.75 |
| **AtlasTrack** | L CgH | -0.0051 | 0.00027 | -23.94 | 1.23 | -0.015 | 0.00039 | -37.78 |
| **AtlasTrack** | R CST | -0.0065 | 0.00053 | -29.78 | 2.33 | -0.014 | 0.00032 | -45.42 |
| **AtlasTrack** | L CST | -0.0078 | 0.00086 | -40.32 | 4.08 | -0.013 | 0.00031 | -42.37 |
| **AtlasTrack** | R ATR | -0.01 | 0.0029 | -47.92 | 12.06 | -0.016 | 0.00036 | -43.56 |
| **AtlasTrack** | L ATR | -0.013 | 0.0036 | -56.81 | 16.63 | -0.015 | 0.00037 | -39.12 |
| **AtlasTrack** | R Unc | -0.0045 | 0.0013 | -29.66 | 6.37 | -0.0096 | 0.0003 | -32.42 |
| **AtlasTrack** | L Unc | -0.0046 | 0.00063 | -33.11 | 3.19 | -0.01 | 0.00029 | -34.35 |
| **AtlasTrack** | R ILF | -0.0051 | 0.00036 | -29.23 | 1.79 | -0.011 | 0.00031 | -33.62 |
| **AtlasTrack** | L ILF | -0.004 | 0.00015 | -23.18 | 0.8 | -0.01 | 0.00035 | -29.34 |
| **AtlasTrack** | R IFO | -0.0053 | 0.0013 | -29.66 | 6.37 | -0.011 | 0.00028 | -39.38 |
| **AtlasTrack** | L IFO | -0.006 | 0.002 | -31.59 | 8.49 | -0.011 | 0.00029 | -38.86 |
| **AtlasTrack** | R SLF | -0.0054 | 0.00047 | -30.41 | 2.26 | -0.013 | 0.00033 | -39.76 |
| **AtlasTrack** | L SLF | -0.005 | -0.00047 | -27.71 | -2.09 | -0.012 | 0.00029 | -41.54 |
| **AtlasTrack** | R SCS | -0.0086 | -0.00034 | -41.19 | -1.48 | -0.019 | 0.00036 | -53.95 |
| **AtlasTrack** | L SCS | -0.0089 | 0.00026 | -41.85 | 1.24 | -0.018 | 0.00034 | -52.77 |
| **AtlasTrack** | R SIFC | -0.0078 | 0.0015 | -39.03 | 6.92 | -0.013 | 0.00033 | -37.43 |
| **AtlasTrack** | L SIFC | -0.0072 | 0.002 | -31.59 | 8.49 | -0.014 | 0.00029 | -48.74 |
| **AtlasTrack** | R IFSFC | -0.0055 | -0.00042 | -24.98 | -1.98 | -0.014 | 0.00035 | -40.24 |
| **AtlasTrack** | L IFSFC | -0.0047 | -0.0003 | -21.49 | -1.43 | -0.012 | 0.00032 | -38.47 |
| **Freesurfer Aseg** | Ca | -0.0052 | 0.0028 | -24.75 | 11.59 | -0.0075 | 0.00038 | -19.47 |
| **Freesurfer Aseg** | Pu | -0.0089 | 0.0059 | -41.85 | 26.74 | -0.023 | 0.00033 | -69.43 |
| **Freesurfer Aseg** | GP | -0.011 | 0.0031 | -50.83 | 13.64 | -0.029 | 0.00033 | -89.75 |
| **Freesurfer Aseg** | Thal | -0.0081 | 0.0026 | -35.67 | 11.03 | -0.019 | 0.00033 | -58.11 |
| **Freesurfer Aseg** | NAcc | -0.0055 | -0.00022 | -25.98 | -0.93 | -0.017 | 0.00039 | -43.55 |
| **Freesurfer Aseg** | VDC | -0.0096 | 0.0026 | -51.15 | 12.61 | -0.022 | 0.00033 | -67.45 |
| **Freesurfer Aseg** | Hipp | -0.0034 | 0.0021 | -15.21 | 13.27 | -0.0033 | 0.00034 | -9.86 |
| **Freesurfer Aseg** | Amy | -0.005 | 0.0011 | -24.38 | 4.33 | -0.012 | 0.00041 | -29.1 |
| **Najdenovska** | A | -0.0074 | 0.0026 | -34.42 | 11.03 | -0.017 | 0.0004 | -40.85 |
| **Najdenovska** | MD | -0.007 | 0.0016 | -32.53 | 7.11 | -0.017 | 0.00037 | -45.99 |
| **Najdenovska** | VA | -0.0073 | -0.00047 | -33.95 | -2.17 | -0.021 | 0.00039 | -53.06 |
| **Najdenovska** | VLD | -0.0075 | 0.00043 | -33.47 | 2.71 | -0.015 | 0.00034 | -43.7 |
| **Najdenovska** | VLV | -0.0081 | 0.0026 | -35.67 | 10.81 | -0.018 | 0.00038 | -46.98 |
| **Najdenovska** | CllpmPUL | -0.0045 | 0.0013 | -19.57 | 5.82 | -0.007 | 0.00035 | -20.06 |
| **Najdenovska** | PUL | -0.0056 | 0.0011 | -27.28 | 5.35 | -0.014 | 0.00034 | -40.16 |
| **Pauli** | SNc | -0.0096 | -0.0058 | -51.15 | -30.34 | -0.027 | 0.00037 | -72.01 |
| **Pauli** | RN | -0.0088 | -0.0033 | -43.57 | -14.19 | -0.023 | 0.00037 | -61.5 |
| **Pauli** | SNr | -0.0094 | -0.0044 | -51.15 | -22.62 | -0.028 | 0.00035 | -79.06 |
| **Pauli** | PBP | -0.0079 | -0.0064 | -37.34 | -31.06 | -0.023 | 0.00041 | -56.75 |
| **Pauli** | HTH | -0.006 | 0.0036 | -29.48 | 14.19 | -0.0029 | 0.00048 | -6.15 |
| **Pauli** | MN | -0.0064 | 0.0022 | -29.42 | 9.04 | 0.00062 | 0.00054 | 1.16 |
| **Pauli** | STH | -0.009 | -0.0044 | -46.1 | -18.22 | -0.024 | 0.0004 | -59.41 |

**Supplementary Table 7. Summary statistics for the FNT age associations.** Statistics include the range of beta coefficients and t statistics within each ROI for the voxelwise analyses and the estimated beta coefficient, SE and t statistic the ROI analyses.

| | | age voxelwise effects | | | | age ROIwise effects | | |
|---|---|---|---|---|---|---|---|---|
| **Atlas** | **ROI** | $min\beta$ | $max\beta$ | $min\ t$ | $max\ t$ | $\beta$ | $SE$ | $t$ |
| **AtlasTrack** | CC | -0.0056 | 0.0037 | -25.86 | 14.6 | -0.012 | 0.00041 | -29.58 |
| **AtlasTrack** | Fmaj | -0.003 | 0.0037 | -15.41 | 14.6 | -0.0066 | 0.00045 | -14.74 |
| **AtlasTrack** | Fmin | -0.0026 | 0.0029 | -10.78 | 14.11 | -0.004 | 0.00048 | -8.42 |
| **AtlasTrack** | R Fx | -0.0031 | 0.0035 | -12.39 | 16.8 | 0.0029 | 0.00047 | 6.14 |
| **AtlasTrack** | L Fx | -0.0025 | 0.0034 | -11.04 | 19.57 | 0.0037 | 0.00045 | 8.26 |
| **AtlasTrack** | R CgC | -0.0038 | -0.0012 | -14.46 | -4.57 | -0.012 | 0.00055 | -21.11 |
| **AtlasTrack** | L CgC | -0.004 | -0.0019 | -14.73 | -7.03 | -0.013 | 0.00058 | -22.4 |
| **AtlasTrack** | R CgH | -0.0035 | -0.00094 | -14.12 | -3.94 | -0.01 | 0.00051 | -19.94 |
| **AtlasTrack** | L CgH | -0.0036 | -0.0011 | -14.67 | -4.22 | -0.011 | 0.00047 | -22.76 |
| **AtlasTrack** | R CST | -0.0042 | 0.0012 | -17.26 | 4.56 | -0.0086 | 0.00046 | -18.81 |
| **AtlasTrack** | L CST | -0.0051 | 0.0012 | -20.18 | 4.81 | -0.009 | 0.00046 | -19.7 |
| **AtlasTrack** | R ATR | -0.0033 | 0.00089 | -13.26 | 3.51 | -0.0037 | 0.0005 | -7.51 |
| **AtlasTrack** | L ATR | -0.0036 | 0.0002 | -16.39 | 0.75 | -0.0077 | 0.0005 | -15.31 |
| **AtlasTrack** | R Unc | -0.0024 | 0.0011 | -10.8 | 4.46 | -0.0055 | 0.00048 | -11.61 |
| **AtlasTrack** | L Unc | -0.003 | 0.00044 | -12.08 | 1.77 | -0.0096 | 0.0005 | -18.98 |
| **AtlasTrack** | R ILF | -0.0035 | -0.00065 | -15.22 | -2.73 | -0.011 | 0.00046 | -24.33 |
| **AtlasTrack** | L ILF | -0.0045 | -0.00011 | -18.86 | -0.47 | -0.012 | 0.00046 | -25.74 |
| **AtlasTrack** | R IFO | -0.0034 | 0.0018 | -13.38 | 8.52 | -0.0091 | 0.00048 | -18.98 |
| **AtlasTrack** | L IFO | -0.0047 | 0.0015 | -19.44 | 8.61 | -0.012 | 0.00047 | -25.87 |
| **AtlasTrack** | R SLF | -0.0043 | 3.60E-05 | -17.94 | 0.15 | -0.015 | 0.00046 | -33.04 |
| **AtlasTrack** | L SLF | -0.0052 | -0.002 | -22.12 | -8.13 | -0.016 | 0.00044 | -36.87 |
| **AtlasTrack** | R SCS | -0.0056 | 0.00037 | -25.56 | 1.5 | -0.012 | 0.00043 | -27.5 |
| **AtlasTrack** | L SCS | -0.0061 | -0.00038 | -25.15 | -1.65 | -0.015 | 0.00045 | -32.63 |
| **AtlasTrack** | R SIFC | -0.0028 | 0.00032 | -10.72 | 1.23 | -0.0065 | 0.00055 | -11.84 |
| **AtlasTrack** | L SIFC | -0.0038 | -0.00046 | -15.03 | -1.87 | -0.0092 | 0.00049 | -18.86 |
| **AtlasTrack** | R IFSFC | -0.004 | -0.00091 | -16.53 | -3.56 | -0.015 | 0.00047 | -32.45 |
| **AtlasTrack** | L IFSFC | -0.0042 | -0.00091 | -16.81 | -3.39 | -0.015 | 0.00047 | -31.92 |
| **Freesurfer Aseg** | Ca | -0.005 | 0.0038 | -22.87 | 16.58 | -0.0091 | 0.0005 | -18.1 |
| **Freesurfer Aseg** | Pu | -0.0045 | 0.0018 | -17.53 | 7.73 | -0.01 | 0.00049 | -21.05 |
| **Freesurfer Aseg** | GP | -0.0027 | 0.0027 | -10.52 | 10.37 | 5.80E-05 | 0.00051 | 0.11 |
| **Freesurfer Aseg** | Thal | -0.0042 | 0.0037 | -16.99 | 17.5 | -0.0054 | 0.00046 | -11.84 |
| **Freesurfer Aseg** | NAcc | -0.0027 | 0.001 | -10.82 | 3.84 | -0.0021 | 0.00055 | -3.76 |
| **Freesurfer Aseg** | VDC | -0.0044 | 0.0012 | -18.05 | 4.56 | -0.0057 | 0.00048 | -11.67 |
| **Freesurfer Aseg** | Hipp | -0.004 | 0.002 | -15.95 | 9.23 | -0.009 | 0.00044 | -20.42 |
| **Freesurfer Aseg** | Amy | -0.0044 | 0.00085 | -17.28 | 3.54 | -0.013 | 0.00047 | -27.48 |
| **Najdenovska** | A | -0.003 | 0.003 | -12.79 | 12.01 | -0.0021 | 0.00048 | -4.47 |
| **Najdenovska** | MD | -0.0034 | 0.0013 | -14.34 | 5.76 | -0.006 | 0.00047 | -12.81 |
| **Najdenovska** | VA | -0.0026 | 0.0033 | -10.97 | 15.07 | -0.0024 | 0.00046 | -5.15 |
| **Najdenovska** | VLD | -0.0042 | 0.0034 | -16.99 | 15.9 | -0.0051 | 0.00046 | -11.11 |
| **Najdenovska** | VLV | -0.0024 | 0.00047 | -9.82 | 1.88 | -0.0044 | 0.00045 | -9.87 |
| **Najdenovska** | CllpmPUL | -0.0038 | 0.0022 | -14.89 | 9.59 | -0.0096 | 0.0005 | -19.3 |
| **Najdenovska** | PUL | -0.0041 | 0.0031 | -16.46 | 12.83 | -0.0059 | 0.00051 | -11.72 |
| **Pauli** | SNc | -0.0018 | -8.60E-05 | -7.14 | -0.34 | -0.0022 | 0.00051 | -4.31 |
| **Pauli** | RN | -0.0019 | 0.00023 | -7.35 | 0.89 | -0.0026 | 0.00051 | -5.01 |
| **Pauli** | SNr | -0.002 | 0.0007 | -7.86 | 2.81 | -0.00072 | 0.00049 | -1.47 |
| **Pauli** | PBP | -0.0012 | 0.00026 | -4.71 | 0.99 | -0.00038 | 0.00054 | -0.7 |
| **Pauli** | HTH | -0.004 | 0.00073 | -15.84 | 3.23 | -0.0092 | 0.00054 | -17.13 |
| **Pauli** | MN | -0.004 | -0.0002 | -17.76 | -0.81 | -0.0095 | 0.00057 | -16.59 |
| **Pauli** | STH | -0.0016 | 0.0011 | -5.99 | 4.34 | 0.00037 | 0.00054 | 0.68 |

**Supplementary Table 8. Summary statistics for the RNI age associations.** Statistics include the range of beta coefficients and t statistics within each ROI for the voxelwise analyses and the estimated beta coefficient, SE and t statistic the ROI analyses.

| | | age voxelwise effects | | | | age ROIwise effects | | |
|---|---|---|---|---|---|---|---|---|
| **Atlas** | **ROI** | $min\beta$ | $max\beta$ | $min\ t$ | $max\ t$ | $\beta$ | $SE$ | $t$ |
| **AtlasTrack** | CC | -0.002 | 0.0063 | -10.32 | 38.31 | 0.012 | 0.0003 | 39.27 |
| **AtlasTrack** | Fmaj | -0.00041 | 0.0059 | -2.1 | 33.88 | 0.015 | 0.00029 | 52.2 |
| **AtlasTrack** | Fmin | -0.002 | 0.0044 | -10.32 | 20.11 | 0.008 | 0.00041 | 19.46 |
| **AtlasTrack** | R Fx | -0.0015 | 0.0053 | -8.92 | 24.18 | 0.0094 | 0.00036 | 26.45 |
| **AtlasTrack** | L Fx | -0.0018 | 0.0053 | -11.45 | 25.32 | 0.0072 | 0.00034 | 20.96 |
| **AtlasTrack** | R CgC | 0.0007 | 0.005 | 3.21 | 24.97 | 0.0099 | 0.00035 | 28.14 |
| **AtlasTrack** | L CgC | -0.00056 | 0.0048 | -2.6 | 23.5 | 0.0086 | 0.00034 | 25.14 |
| **AtlasTrack** | R CgH | 0.0038 | 0.0067 | 17.95 | 33.37 | 0.019 | 0.00034 | 56.4 |
| **AtlasTrack** | L CgH | 0.0041 | 0.0064 | 18.08 | 31.91 | 0.019 | 0.00035 | 53.79 |
| **AtlasTrack** | R CST | -0.0011 | 0.0068 | -5.21 | 32.26 | 0.015 | 0.00034 | 43.62 |
| **AtlasTrack** | L CST | -0.00092 | 0.01 | -4.47 | 45.05 | 0.016 | 0.00031 | 53.69 |
| **AtlasTrack** | R ATR | -0.00043 | 0.01 | -2.06 | 49.18 | 0.016 | 0.00036 | 44.24 |
| **AtlasTrack** | L ATR | -0.00087 | 0.011 | -4.02 | 50.32 | 0.016 | 0.00034 | 48.31 |
| **AtlasTrack** | R Unc | 0.0018 | 0.0071 | 9.36 | 30.27 | 0.017 | 0.00035 | 48.12 |
| **AtlasTrack** | L Unc | 0.0022 | 0.0066 | 10.8 | 30.9 | 0.017 | 0.0004 | 42.32 |
| **AtlasTrack** | R ILF | 0.0011 | 0.0064 | 6.56 | 34.58 | 0.016 | 0.00032 | 50.71 |
| **AtlasTrack** | L ILF | 4.70E-05 | 0.0059 | 0.28 | 29.46 | 0.013 | 0.00033 | 40.01 |
| **AtlasTrack** | R IFO | 0.00024 | 0.0075 | 1.15 | 33.64 | 0.015 | 0.00032 | 47.31 |
| **AtlasTrack** | L IFO | -0.00027 | 0.0086 | -1.58 | 36.13 | 0.013 | 0.00031 | 42.85 |
| **AtlasTrack** | R SLF | -0.00093 | 0.0076 | -5.66 | 37.41 | 0.017 | 0.0003 | 56.41 |
| **AtlasTrack** | L SLF | 0.00021 | 0.0058 | 1.41 | 32.08 | 0.017 | 0.00032 | 53.56 |
| **AtlasTrack** | R SCS | 0.0012 | 0.0097 | 6.52 | 43.66 | 0.019 | 0.00032 | 60.25 |
| **AtlasTrack** | L SCS | 0.0012 | 0.01 | 7.56 | 44.46 | 0.019 | 0.00033 | 58.48 |
| **AtlasTrack** | R SIFC | 0.00069 | 0.012 | 3.2 | 58.57 | 0.018 | 0.00042 | 42.88 |
| **AtlasTrack** | L SIFC | 0.0013 | 0.012 | 5.92 | 53.48 | 0.018 | 0.00039 | 46.63 |
| **AtlasTrack** | R IFSFC | 0.0013 | 0.0051 | 6.84 | 31.1 | 0.018 | 0.00035 | 49.58 |
| **AtlasTrack** | L IFSFC | 0.0012 | 0.0051 | 7.56 | 28.95 | 0.015 | 0.00034 | 45.99 |
| **Freesurfer Aseg** | Ca | -0.0026 | 0.0085 | -14.34 | 36.51 | 0.022 | 0.0004 | 56.31 |
| **Freesurfer Aseg** | Pu | 0.0019 | 0.01 | 7.82 | 44.88 | 0.028 | 0.00038 | 74.78 |
| **Freesurfer Aseg** | GP | -3.70E-05 | 0.012 | -0.15 | 61.19 | 0.029 | 0.00032 | 89.08 |
| **Freesurfer Aseg** | Thal | -0.0034 | 0.0088 | -17.16 | 40.04 | 0.019 | 0.00032 | 60.16 |
| **Freesurfer Aseg** | NAcc | 0.0052 | 0.009 | 23.55 | 41.21 | 0.023 | 0.00037 | 61.74 |
| **Freesurfer Aseg** | VDC | -0.00025 | 0.012 | -1.23 | 57.42 | 0.023 | 0.00034 | 69.18 |
| **Freesurfer Aseg** | Hipp | 0.00012 | 0.0067 | 0.59 | 29.23 | 0.019 | 0.00036 | 51.67 |
| **Freesurfer Aseg** | Amy | 0.0025 | 0.0076 | 10.59 | 34.99 | 0.021 | 0.0004 | 52.42 |
| **Najdenovska** | A | -0.0014 | 0.0081 | -5.91 | 37.23 | 0.016 | 0.00036 | 45.05 |
| **Najdenovska** | MD | -0.00076 | 0.0088 | -3.17 | 39.69 | 0.019 | 0.00035 | 53.74 |
| **Najdenovska** | VA | -0.002 | 0.0088 | -9.83 | 40.04 | 0.02 | 0.00037 | 53.79 |
| **Najdenovska** | VLD | -0.0023 | 0.0082 | -11.95 | 37.47 | 0.018 | 0.00035 | 51.92 |
| **Najdenovska** | VLV | 0.00032 | 0.0086 | 1.42 | 39.67 | 0.021 | 0.00034 | 62.02 |
| **Najdenovska** | CllpmPUL | 0.0016 | 0.0065 | 6.83 | 30.96 | 0.016 | 0.00036 | 45.4 |
| **Najdenovska** | PUL | 0.0011 | 0.0069 | 5.4 | 31.36 | 0.016 | 0.00033 | 48.95 |
| **Pauli** | SNc | 0.0051 | 0.012 | 24.13 | 54.49 | 0.028 | 0.00038 | 74.42 |
| **Pauli** | RN | 0.0021 | 0.0088 | 10.32 | 40.65 | 0.022 | 0.00036 | 59.22 |
| **Pauli** | SNr | 0.0055 | 0.012 | 26.49 | 57.42 | 0.03 | 0.00034 | 90.17 |
| **Pauli** | PBP | 0.0064 | 0.0081 | 27.68 | 34.98 | 0.024 | 0.00042 | 57.85 |
| **Pauli** | HTH | 0.0018 | 0.0085 | 7.93 | 38.5 | 0.02 | 0.00039 | 52.23 |
| **Pauli** | MN | 0.0047 | 0.0073 | 21.79 | 34.31 | 0.017 | 0.00041 | 40.44 |
| **Pauli** | STH | 0.0027 | 0.01 | 11.21 | 46.49 | 0.024 | 0.0004 | 59.78 |

**Supplementary Table 9. Summary statistics for the RND age associations.** Statistics include the range of beta coefficients and t statistics within each ROI for the voxelwise analyses and the estimated beta coefficient, SE and t statistic the ROI analyses.

| | | age voxelwise effects | | | | age ROIwise effects | | |
|---|---|---|---|---|---|---|---|---|
| **Atlas** | **ROI** | $min\beta$ | $max\beta$ | $min\ t$ | $max\ t$ | $\beta$ | $SE$ | $t$ |
| **AtlasTrack** | CC | -0.0054 | 0.0045 | -30.91 | 30.38 | 0.006 | 0.00021 | 27.8 |
| **AtlasTrack** | Fmaj | -0.0054 | 0.0025 | -30.91 | 19.44 | 0.0035 | 0.00024 | 14.61 |
| **AtlasTrack** | Fmin | -0.003 | 0.0022 | -13.01 | 11.83 | 0.0012 | 0.00023 | 4.94 |
| **AtlasTrack** | R Fx | -0.0019 | 0.0029 | -10.11 | 17.26 | 0.0052 | 0.00025 | 21 |
| **AtlasTrack** | L Fx | -0.0018 | 0.003 | -8.84 | 13.93 | 0.0041 | 0.00027 | 15.13 |
| **AtlasTrack** | R CgC | -0.0012 | 0.004 | -7.87 | 24.17 | 0.0072 | 0.00029 | 24.93 |
| **AtlasTrack** | L CgC | -0.0027 | 0.0042 | -13.52 | 25.82 | 0.0066 | 0.00033 | 19.61 |
| **AtlasTrack** | R CgH | -0.00082 | 0.0033 | -4.67 | 16.57 | 0.01 | 0.0003 | 34.41 |
| **AtlasTrack** | L CgH | -0.0014 | 0.0037 | -7.16 | 19.28 | 0.011 | 0.00028 | 38.02 |
| **AtlasTrack** | R CST | -0.0037 | 0.0029 | -18.04 | 15.65 | 0.0046 | 0.00025 | 18.5 |
| **AtlasTrack** | L CST | -0.0058 | 0.0047 | -26.38 | 21.17 | 0.003 | 0.00024 | 12.27 |
| **AtlasTrack** | R ATR | -0.0043 | 0.0062 | -19.32 | 28.23 | 0.0093 | 0.00024 | 38.86 |
| **AtlasTrack** | L ATR | -0.0064 | 0.0068 | -32.68 | 29.29 | 0.0063 | 0.00027 | 23.07 |
| **AtlasTrack** | R Unc | -0.0037 | 0.0024 | -16.25 | 16.93 | 0.0049 | 0.00023 | 21.93 |
| **AtlasTrack** | L Unc | -0.0017 | 0.0027 | -9.11 | 18.09 | 0.0073 | 0.00022 | 33.32 |
| **AtlasTrack** | R ILF | -0.0024 | 0.0027 | -15.78 | 17.98 | 0.006 | 0.00025 | 24.28 |
| **AtlasTrack** | L ILF | -0.0023 | 0.0033 | -12.26 | 20.03 | 0.008 | 0.00028 | 28.83 |
| **AtlasTrack** | R IFO | -0.0037 | 0.0031 | -16.55 | 18.93 | 0.0076 | 0.00021 | 35.89 |
| **AtlasTrack** | L IFO | -0.0038 | 0.0041 | -16 | 21 | 0.0084 | 0.00024 | 35.5 |
| **AtlasTrack** | R SLF | -0.0028 | 0.0034 | -12.8 | 20.13 | 0.007 | 0.00023 | 30.12 |
| **AtlasTrack** | L SLF | -0.0016 | 0.0029 | -9.26 | 16.96 | 0.0059 | 0.00022 | 26.29 |
| **AtlasTrack** | R SCS | -0.0025 | 0.0061 | -13.71 | 28.98 | 0.013 | 0.00027 | 49.6 |
| **AtlasTrack** | L SCS | -0.0038 | 0.0072 | -17.08 | 33.52 | 0.014 | 0.00026 | 52.89 |
| **AtlasTrack** | R SIFC | -0.0058 | 0.0047 | -24.67 | 22.38 | 0.0072 | 0.00031 | 23.28 |
| **AtlasTrack** | L SIFC | -0.0038 | 0.004 | -16 | 19.05 | 0.0099 | 0.00027 | 36.6 |
| **AtlasTrack** | R IFSFC | -0.00082 | 0.0028 | -4.24 | 14.96 | 0.0081 | 0.00024 | 33.79 |
| **AtlasTrack** | L IFSFC | -0.001 | 0.0026 | -5.99 | 15.84 | 0.0082 | 0.00024 | 33.73 |
| **Freesurfer Aseg** | Ca | -0.0048 | 0.0059 | -22.49 | 27.19 | 0.0069 | 0.00024 | 28.41 |
| **Freesurfer Aseg** | Pu | -0.0086 | 0.0072 | -37.94 | 33.52 | 0.012 | 0.00025 | 47.92 |
| **Freesurfer Aseg** | GP | -0.0065 | 0.0071 | -27.49 | 30.41 | 0.0065 | 0.0003 | 21.48 |
| **Freesurfer Aseg** | Thal | -0.0043 | 0.0062 | -20.23 | 28.23 | 0.013 | 0.00024 | 56.32 |
| **Freesurfer Aseg** | NAcc | -0.0011 | 0.0045 | -4.81 | 19.57 | 0.012 | 0.00035 | 34.2 |
| **Freesurfer Aseg** | VDC | -0.0043 | 0.0075 | -19.64 | 33.67 | 0.016 | 0.00026 | 62.55 |
| **Freesurfer Aseg** | Hipp | -0.002 | 0.0035 | -10.83 | 17.41 | 0.0057 | 0.00024 | 23.96 |
| **Freesurfer Aseg** | Amy | -0.0023 | 0.0048 | -11.58 | 21.45 | 0.0079 | 0.00032 | 24.72 |
| **Najdenovska** | A | -0.0029 | 0.0062 | -14.49 | 28.23 | 0.011 | 0.0003 | 36.32 |
| **Najdenovska** | MD | -0.0025 | 0.0049 | -11.55 | 20.25 | 0.013 | 0.00035 | 36.71 |
| **Najdenovska** | VA | -0.0027 | 0.0062 | -12.55 | 27.89 | 0.017 | 0.00031 | 53.54 |
| **Najdenovska** | VLD | -0.0024 | 0.005 | -11.15 | 21.62 | 0.0095 | 0.00028 | 33.8 |
| **Najdenovska** | VLV | -0.0044 | 0.0042 | -20.23 | 17.24 | 0.00049 | 0.00025 | 1.95 |
| **Najdenovska** | CllpmPUL | -0.0025 | 0.0038 | -12.08 | 17.74 | 0.002 | 0.00028 | 7.19 |
| **Najdenovska** | PUL | -0.0039 | 0.004 | -17.08 | 20.32 | 0.0044 | 0.00031 | 14.47 |
| **Pauli** | SNc | 0.0028 | 0.0075 | 14.42 | 33.42 | 0.021 | 0.00035 | 61.92 |
| **Pauli** | RN | -0.00065 | 0.0067 | -3.06 | 26.75 | 0.017 | 0.00037 | 46.89 |
| **Pauli** | SNr | 0.00011 | 0.0075 | 0.57 | 33.67 | 0.021 | 0.00036 | 57.62 |
| **Pauli** | PBP | 0.0016 | 0.0043 | 8.54 | 18.91 | 0.016 | 0.00036 | 43.69 |
| **Pauli** | HTH | -0.003 | 0.0052 | -15.84 | 22.19 | 0.0058 | 0.00032 | 18.53 |
| **Pauli** | MN | 0.00022 | 0.0055 | 1.06 | 22.27 | 0.013 | 0.00055 | 24.12 |
| **Pauli** | STH | -0.00067 | 0.0038 | -3.27 | 19.59 | 0.011 | 0.00036 | 30.01 |

**Supplementary Table 10. Summary statistics for the RDF age associations.** Statistics include the range of beta coefficients and t statistics within each ROI for the voxelwise analyses and the estimated beta coefficient, SE and t statistic the ROI analyses.

| | | age voxelwise effects | | | | age ROIwise effects | | |
|---|---|---|---|---|---|---|---|---|
| **Atlas** | **ROI** | $min\beta$ | $max\beta$ | $min\ t$ | $max\ t$ | $\beta$ | $SE$ | $t$ |
| **AtlasTrack** | CC | -0.0053 | 0.0031 | -35.31 | 16.86 | -0.0092 | 0.00026 | -35.2 |
| **AtlasTrack** | Fmaj | -0.0053 | 0.0012 | -35.31 | 7.67 | -0.013 | 0.00026 | -48.32 |
| **AtlasTrack** | Fmin | -0.0036 | 0.0011 | -16.6 | 5.26 | -0.0071 | 0.00028 | -25.25 |
| **AtlasTrack** | R Fx | -0.0029 | 0.0025 | -15.58 | 14.6 | -0.0031 | 0.00035 | -8.99 |
| **AtlasTrack** | L Fx | -0.0033 | 0.0018 | -17.47 | 9.61 | -0.0022 | 0.00033 | -6.72 |
| **AtlasTrack** | R CgC | -0.0026 | 0.0021 | -15.26 | 11.96 | -0.0035 | 0.00033 | -10.43 |
| **AtlasTrack** | L CgC | -0.0038 | 0.0024 | -20.03 | 14.4 | -0.0049 | 0.00033 | -14.66 |
| **AtlasTrack** | R CgH | -0.0029 | 0.00078 | -17.39 | 4.05 | -0.0056 | 0.00033 | -17.14 |
| **AtlasTrack** | L CgH | -0.0034 | 0.00095 | -19.88 | 5.41 | -0.0054 | 0.00032 | -16.73 |
| **AtlasTrack** | R CST | -0.0049 | 0.0011 | -26.6 | 5.99 | -0.0091 | 0.00029 | -31.99 |
| **AtlasTrack** | L CST | -0.0071 | 0.00077 | -35.13 | 4.15 | -0.011 | 0.00028 | -40.11 |
| **AtlasTrack** | R ATR | -0.0066 | 0.0032 | -30.2 | 15.64 | -0.0067 | 0.0003 | -21.96 |
| **AtlasTrack** | L ATR | -0.0076 | 0.0037 | -39.66 | 16.94 | -0.01 | 0.00031 | -31.77 |
| **AtlasTrack** | R Unc | -0.0066 | -4.90E-05 | -27.55 | -0.31 | -0.014 | 0.00032 | -44.15 |
| **AtlasTrack** | L Unc | -0.0047 | 0.00048 | -23.79 | 3.09 | -0.009 | 0.00031 | -29.62 |
| **AtlasTrack** | R ILF | -0.0045 | 0.00069 | -27.99 | 4.43 | -0.013 | 0.00025 | -50.55 |
| **AtlasTrack** | L ILF | -0.0043 | 0.0013 | -24.51 | 7.73 | -0.0087 | 0.0003 | -29.23 |
| **AtlasTrack** | R IFO | -0.0066 | 0.0011 | -28.54 | 7.06 | -0.012 | 0.00029 | -40.52 |
| **AtlasTrack** | L IFO | -0.0068 | 0.0013 | -29.98 | 7.73 | -0.0092 | 0.00029 | -31.8 |
| **AtlasTrack** | R SLF | -0.0054 | 0.0016 | -24.2 | 9.95 | -0.011 | 0.00028 | -38.65 |
| **AtlasTrack** | L SLF | -0.0045 | 0.001 | -23.08 | 5.72 | -0.011 | 0.00029 | -39.69 |
| **AtlasTrack** | R SCS | -0.0051 | 0.0035 | -24.96 | 15.41 | -0.0067 | 0.0003 | -22.33 |
| **AtlasTrack** | L SCS | -0.0056 | 0.0043 | -28.34 | 20.55 | -0.008 | 0.00031 | -25.83 |
| **AtlasTrack** | R SIFC | -0.007 | 0.0021 | -34.67 | 9.62 | -0.013 | 0.00042 | -31.08 |
| **AtlasTrack** | L SIFC | -0.007 | 0.0011 | -29.99 | 5.11 | -0.011 | 0.0004 | -26.33 |
| **AtlasTrack** | R IFSFC | -0.0028 | 0.00021 | -16.14 | 1.11 | -0.0088 | 0.0003 | -28.97 |
| **AtlasTrack** | L IFSFC | -0.0028 | 0.00039 | -15.45 | 2.25 | -0.0084 | 0.00031 | -26.77 |
| **Freesurfer Aseg** | Ca | -0.0066 | 0.0042 | -30.2 | 18.67 | -0.0034 | 0.00025 | -13.37 |
| **Freesurfer Aseg** | Pu | -0.0094 | 0.0043 | -42.97 | 20.55 | -0.0068 | 0.00029 | -23.58 |
| **Freesurfer Aseg** | GP | -0.0074 | 0.00042 | -39.87 | 1.91 | -0.017 | 0.00026 | -63.39 |
| **Freesurfer Aseg** | Thal | -0.005 | 0.004 | -30.47 | 18.4 | -0.0067 | 0.0003 | -22.46 |
| **Freesurfer Aseg** | NAcc | -0.0042 | 0.00072 | -20.07 | 3.07 | -0.0042 | 0.00035 | -11.77 |
| **Freesurfer Aseg** | VDC | -0.0065 | 0.0045 | -37.55 | 17.99 | -0.012 | 0.00034 | -35.55 |
| **Freesurfer Aseg** | Hipp | -0.0039 | 0.0014 | -18.81 | 6.71 | -0.0076 | 0.0003 | -25.73 |
| **Freesurfer Aseg** | Amy | -0.005 | 0.0021 | -25.01 | 9.59 | -0.0068 | 0.00033 | -20.42 |
| **Najdenovska** | A | -0.0039 | 0.0032 | -20.19 | 15.64 | -0.0038 | 0.00033 | -11.47 |
| **Najdenovska** | MD | -0.0046 | 0.0022 | -23.43 | 9.83 | -0.0027 | 0.00029 | -9.42 |
| **Najdenovska** | VA | -0.0055 | 0.004 | -28.77 | 18.4 | -0.00045 | 0.00032 | -1.39 |
| **Najdenovska** | VLD | -0.0048 | 0.0031 | -24.32 | 13.64 | -0.0043 | 0.00032 | -13.66 |
| **Najdenovska** | VLV | -0.0069 | -0.00066 | -34.87 | -2.94 | -0.013 | 0.00028 | -47 |
| **Najdenovska** | CllpmPUL | -0.0044 | 0.0026 | -22.73 | 11.8 | -0.0068 | 0.00027 | -24.84 |
| **Najdenovska** | PUL | -0.0049 | 0.00093 | -30.47 | 5.6 | -0.0083 | 0.0003 | -27.88 |
| **Pauli** | SNc | -0.0021 | 0.00019 | -9.51 | 1.01 | -0.0029 | 0.00034 | -8.52 |
| **Pauli** | RN | -0.0029 | 0.0045 | -14.1 | 17.99 | -0.0019 | 0.00038 | -5.15 |
| **Pauli** | SNr | -0.0065 | -8.60E-05 | -35.55 | -0.44 | -0.013 | 0.00036 | -35.29 |
| **Pauli** | PBP | -0.0026 | -0.00027 | -16.22 | -1.4 | -0.0054 | 0.0004 | -13.58 |
| **Pauli** | HTH | -0.0054 | 0.0012 | -29.03 | 6.24 | -0.012 | 0.00031 | -39.58 |
| **Pauli** | MN | -0.0025 | 0.00036 | -12.59 | 1.54 | -0.0033 | 0.00044 | -7.44 |
| **Pauli** | STH | -0.0046 | 0.00016 | -22.64 | 0.75 | -0.0094 | 0.00038 | -24.66 |

**Supplementary Table 11. Summary statistics for the MD age associations.** Statistics include the range of beta coefficients and t statistics within each ROI for the voxelwise analyses and the estimated beta coefficient, SE and t statistic the ROI analyses.

| | | age voxelwise effects | | | | age ROIwise effects | | |
|---|---|---|---|---|---|---|---|---|
| **Atlas** | **ROI** | $min\beta$ | $max\beta$ | $min\ t$ | $max\ t$ | $\beta$ | $SE$ | $t$ |
| **AtlasTrack** | CC | -0.0075 | 0.0025 | -42.31 | 13.2 | -0.0084 | 0.00027 | -31.15 |
| **AtlasTrack** | Fmaj | -0.0066 | 0.00019 | -37.8 | 1.21 | -0.0096 | 0.00024 | -39.8 |
| **AtlasTrack** | Fmin | -0.0051 | 0.0018 | -27.08 | 9.14 | -0.0051 | 0.00036 | -14.21 |
| **AtlasTrack** | R Fx | -0.0054 | 0.0009 | -31.76 | 5.38 | -0.0069 | 0.00024 | -28.71 |
| **AtlasTrack** | L Fx | -0.0051 | 0.00062 | -31.88 | 3.8 | -0.0055 | 0.00024 | -23 |
| **AtlasTrack** | R CgC | -0.0058 | -0.002 | -28.43 | -10.44 | -0.011 | 0.00032 | -35.45 |
| **AtlasTrack** | L CgC | -0.0054 | -0.0013 | -26.53 | -6.11 | -0.0099 | 0.00032 | -31.32 |
| **AtlasTrack** | R CgH | -0.0076 | -0.0037 | -40.11 | -20.35 | -0.017 | 0.0003 | -57.89 |
| **AtlasTrack** | L CgH | -0.007 | -0.0041 | -38.02 | -20.12 | -0.017 | 0.00031 | -55.73 |
| **AtlasTrack** | R CST | -0.0075 | -0.0009 | -42.81 | -5.38 | -0.014 | 0.00027 | -51.99 |
| **AtlasTrack** | L CST | -0.0095 | -0.00046 | -56.49 | -2.77 | -0.015 | 0.00027 | -55.47 |
| **AtlasTrack** | R ATR | -0.0098 | -0.00022 | -57.12 | -1.06 | -0.013 | 0.00028 | -48.76 |
| **AtlasTrack** | L ATR | -0.011 | -0.0006 | -60.37 | -3.08 | -0.014 | 0.00026 | -53.93 |
| **AtlasTrack** | R Unc | -0.0071 | -0.0013 | -34.59 | -7.29 | -0.014 | 0.0003 | -47.28 |
| **AtlasTrack** | L Unc | -0.007 | -0.0021 | -36.62 | -12.01 | -0.015 | 0.00034 | -43.3 |
| **AtlasTrack** | R ILF | -0.007 | -0.0019 | -41.1 | -11.42 | -0.014 | 0.00031 | -46.19 |
| **AtlasTrack** | L ILF | -0.0066 | -0.00079 | -35.04 | -4.56 | -0.012 | 0.0003 | -39.36 |
| **AtlasTrack** | R IFO | -0.0072 | -0.00039 | -37.07 | -2.13 | -0.013 | 0.0003 | -44.4 |
| **AtlasTrack** | L IFO | -0.0086 | -0.00026 | -39.64 | -1.44 | -0.012 | 0.00027 | -44.04 |
| **AtlasTrack** | R SLF | -0.0084 | -0.003 | -45.27 | -14.6 | -0.018 | 0.00029 | -63.18 |
| **AtlasTrack** | L SLF | -0.0079 | -0.0033 | -43.72 | -15.83 | -0.018 | 0.00031 | -57.53 |
| **AtlasTrack** | R SCS | -0.0092 | -0.0024 | -46.1 | -11.75 | -0.02 | 0.00027 | -71.32 |
| **AtlasTrack** | L SCS | -0.0095 | -0.0022 | -49.68 | -10.28 | -0.02 | 0.00027 | -72.11 |
| **AtlasTrack** | R SIFC | -0.0098 | -0.0011 | -56.52 | -5.25 | -0.016 | 0.00036 | -44.44 |
| **AtlasTrack** | L SIFC | -0.0098 | -0.002 | -52.96 | -9.24 | -0.018 | 0.00034 | -51.82 |
| **AtlasTrack** | R IFSFC | -0.0071 | -0.0025 | -37.98 | -14.48 | -0.018 | 0.00032 | -56.67 |
| **AtlasTrack** | L IFSFC | -0.0065 | -0.0029 | -35.99 | -16.98 | -0.016 | 0.00032 | -51.04 |
| **Freesurfer Aseg** | Ca | -0.0096 | 0.0033 | -44.65 | 15.65 | -0.02 | 0.00033 | -60.16 |
| **Freesurfer Aseg** | Pu | -0.01 | -0.0019 | -50.29 | -8.4 | -0.024 | 0.00031 | -78.11 |
| **Freesurfer Aseg** | GP | -0.01 | 0.00015 | -65.54 | 0.65 | -0.024 | 0.00026 | -91.06 |
| **Freesurfer Aseg** | Thal | -0.0086 | 0.0029 | -49.56 | 14.63 | -0.015 | 0.00025 | -59.93 |
| **Freesurfer Aseg** | NAcc | -0.0078 | -0.0042 | -41.42 | -25.99 | -0.02 | 0.00033 | -60.23 |
| **Freesurfer Aseg** | VDC | -0.011 | -0.00027 | -67.17 | -1.35 | -0.02 | 0.00025 | -80.48 |
| **Freesurfer Aseg** | Hipp | -0.0067 | -0.00035 | -32.98 | -2.1 | -0.015 | 0.00028 | -53.02 |
| **Freesurfer Aseg** | Amy | -0.0092 | -0.0023 | -43.48 | -11.62 | -0.018 | 0.00035 | -53.01 |
| **Najdenovska** | A | -0.0076 | 0.00087 | -44.8 | 4.44 | -0.013 | 0.00027 | -46.72 |
| **Najdenovska** | MD | -0.0081 | 9.80E-05 | -47.89 | 0.54 | -0.015 | 0.00026 | -57.84 |
| **Najdenovska** | VA | -0.0076 | 0.0013 | -44.97 | 7.18 | -0.016 | 0.00028 | -57.45 |
| **Najdenovska** | VLD | -0.0078 | 0.0018 | -43.84 | 10.81 | -0.015 | 0.00027 | -56.6 |
| **Najdenovska** | VLV | -0.0086 | -0.00047 | -49.56 | -2.3 | -0.018 | 0.00026 | -68.18 |
| **Najdenovska** | CllpmPUL | -0.006 | -0.0013 | -38.99 | -8.27 | -0.013 | 0.00026 | -51.3 |
| **Najdenovska** | PUL | -0.0068 | -0.0012 | -39.94 | -7.9 | -0.013 | 0.00024 | -54.56 |
| **Pauli** | SNc | -0.01 | -0.0067 | -64.58 | -36 | -0.024 | 0.00029 | -84.11 |
| **Pauli** | RN | -0.0099 | -0.0046 | -53.07 | -20.84 | -0.021 | 0.00032 | -66.28 |
| **Pauli** | SNr | -0.011 | -0.0063 | -67.17 | -38.86 | -0.025 | 0.00028 | -89.91 |
| **Pauli** | PBP | -0.0082 | -0.007 | -45.8 | -38.64 | -0.021 | 0.00032 | -64.28 |
| **Pauli** | HTH | -0.007 | -0.0016 | -43.44 | -7.76 | -0.015 | 0.00031 | -47.36 |
| **Pauli** | MN | -0.0073 | -0.0044 | -45.74 | -26.95 | -0.013 | 0.00033 | -40.46 |
| **Pauli** | STH | -0.01 | -0.0043 | -57.44 | -21.59 | -0.022 | 0.00031 | -71.11 |

**Supplementary Table 12. Summary statistics for the FA age associations.** Statistics include the range of beta coefficients and t statistics within each ROI for the voxelwise analyses and the estimated beta coefficient, SE and t statistic the ROI analyses.

| | | age voxelwise effects | | | | age ROIwise effects | | |
|---|---|---|---|---|---|---|---|---|
| **Atlas** | **ROI** | $min\beta$ | $max\beta$ | $min\ t$ | $max\ t$ | $\beta$ | $SE$ | $t$ |
| **AtlasTrack** | CC | -0.0035 | 0.0047 | -19.08 | 24.73 | 0.0061 | 0.00026 | 23.59 |
| **AtlasTrack** | Fmaj | -0.0035 | 0.0031 | -19.08 | 21.29 | 0.0057 | 0.00025 | 23.12 |
| **AtlasTrack** | Fmin | -0.0028 | 0.0019 | -12.58 | 11.53 | 0.00034 | 0.00028 | 1.22 |
| **AtlasTrack** | R Fx | -0.0018 | 0.0023 | -10.94 | 12.44 | 0.0039 | 0.0003 | 13.14 |
| **AtlasTrack** | L Fx | -0.0022 | 0.0019 | -11.76 | 9.29 | 0.0039 | 0.0003 | 12.75 |
| **AtlasTrack** | R CgC | -0.001 | 0.0038 | -5.65 | 23.75 | 0.0082 | 0.00031 | 26.67 |
| **AtlasTrack** | L CgC | -0.0027 | 0.0043 | -12.2 | 24.42 | 0.0076 | 0.00033 | 22.83 |
| **AtlasTrack** | R CgH | -0.0011 | 0.0027 | -6.38 | 14.27 | 0.0082 | 0.00031 | 26.96 |
| **AtlasTrack** | L CgH | -0.0016 | 0.0032 | -9.45 | 16.54 | 0.0089 | 0.00032 | 28.11 |
| **AtlasTrack** | R CST | -0.0025 | 0.0031 | -12.61 | 12.97 | 0.0043 | 0.00027 | 15.78 |
| **AtlasTrack** | L CST | -0.0046 | 0.0035 | -20.44 | 15.73 | 0.0029 | 0.00027 | 10.77 |
| **AtlasTrack** | R ATR | -0.0046 | 0.0042 | -20.74 | 20.29 | 0.0053 | 0.00026 | 20.16 |
| **AtlasTrack** | L ATR | -0.0061 | 0.0049 | -28.82 | 25.23 | 0.0031 | 0.00026 | 11.6 |
| **AtlasTrack** | R Unc | -0.004 | 0.0017 | -17.92 | 11.62 | 0.0018 | 0.00031 | 5.93 |
| **AtlasTrack** | L Unc | -0.002 | 0.0023 | -10.91 | 15.42 | 0.0051 | 0.00028 | 18.41 |
| **AtlasTrack** | R ILF | -0.0021 | 0.0023 | -11.17 | 13.27 | 0.0043 | 0.00026 | 16.35 |
| **AtlasTrack** | L ILF | -0.0022 | 0.0029 | -11.48 | 16.83 | 0.006 | 0.00026 | 23.52 |
| **AtlasTrack** | R IFO | -0.004 | 0.0025 | -17.92 | 15.47 | 0.0066 | 0.00029 | 23.03 |
| **AtlasTrack** | L IFO | -0.0044 | 0.003 | -17.75 | 17.14 | 0.0069 | 0.00028 | 25.03 |
| **AtlasTrack** | R SLF | -0.0027 | 0.003 | -12.57 | 16.89 | 0.0075 | 0.00024 | 31.49 |
| **AtlasTrack** | L SLF | -0.0021 | 0.0029 | -11.04 | 15.79 | 0.0064 | 0.00026 | 24.86 |
| **AtlasTrack** | R SCS | -0.0025 | 0.0047 | -11.74 | 21.17 | 0.0069 | 0.00026 | 26.52 |
| **AtlasTrack** | L SCS | -0.0035 | 0.0058 | -16.44 | 25.95 | 0.0066 | 0.00027 | 24.39 |
| **AtlasTrack** | R SIFC | -0.005 | 0.0039 | -24.8 | 17.23 | 0.0046 | 0.00033 | 13.95 |
| **AtlasTrack** | L SIFC | -0.0044 | 0.003 | -17.75 | 12.52 | 0.0072 | 0.00029 | 24.67 |
| **AtlasTrack** | R IFSFC | -0.00096 | 0.0014 | -5.48 | 8.22 | 0.0052 | 0.00026 | 20.12 |
| **AtlasTrack** | L IFSFC | -0.00088 | 0.0017 | -4.78 | 9.79 | 0.0053 | 0.00029 | 18.42 |
| **Freesurfer Aseg** | Ca | -0.0051 | 0.0045 | -22.71 | 21.21 | 0.0045 | 0.00029 | 15.33 |
| **Freesurfer Aseg** | Pu | -0.0083 | 0.0058 | -37.08 | 25.95 | 0.0064 | 0.00028 | 22.9 |
| **Freesurfer Aseg** | GP | -0.006 | 0.006 | -26.98 | 25.33 | 0.0037 | 0.00029 | 12.69 |
| **Freesurfer Aseg** | Thal | -0.0035 | 0.0043 | -17.9 | 20.32 | 0.0061 | 0.00025 | 24.22 |
| **Freesurfer Aseg** | NAcc | -0.0014 | 0.0033 | -6.67 | 14.53 | 0.0064 | 0.00034 | 18.99 |
| **Freesurfer Aseg** | VDC | -0.0039 | 0.0054 | -19.38 | 23.18 | 0.01 | 0.0003 | 34.39 |
| **Freesurfer Aseg** | Hipp | -0.0019 | 0.003 | -10.27 | 14.31 | 0.0029 | 0.00026 | 11.23 |
| **Freesurfer Aseg** | Amy | -0.0029 | 0.0038 | -13.59 | 16.52 | 0.0022 | 0.00028 | 7.87 |
| **Najdenovska** | A | -0.0023 | 0.0039 | -12.23 | 18.35 | 0.0055 | 0.00029 | 19.18 |
| **Najdenovska** | MD | -0.0028 | 0.0041 | -13.86 | 16.97 | 0.0068 | 0.00028 | 24.59 |
| **Najdenovska** | VA | -0.0022 | 0.0043 | -9.97 | 20.32 | 0.011 | 0.00028 | 39.12 |
| **Najdenovska** | VLD | -0.0023 | 0.0033 | -11.01 | 15.86 | 0.0057 | 0.00027 | 20.96 |
| **Najdenovska** | VLV | -0.0035 | 0.002 | -17.2 | 8.44 | -0.0033 | 0.00024 | -13.86 |
| **Najdenovska** | CllpmPUL | -0.0028 | 0.0034 | -14.27 | 14.72 | -0.00053 | 0.00023 | -2.29 |
| **Najdenovska** | PUL | -0.0034 | 0.0024 | -17.9 | 12.74 | 0.00058 | 0.00029 | 1.99 |
| **Pauli** | SNc | 0.0023 | 0.0043 | 12.32 | 18.72 | 0.015 | 0.00035 | 43.29 |
| **Pauli** | RN | -0.00028 | 0.0054 | -1.39 | 23.18 | 0.015 | 0.00038 | 38.85 |
| **Pauli** | SNr | -0.0003 | 0.0046 | -1.49 | 20.08 | 0.012 | 0.00036 | 33.9 |
| **Pauli** | PBP | 0.00086 | 0.0032 | 4.56 | 16.16 | 0.012 | 0.00039 | 30.83 |
| **Pauli** | HTH | -0.0033 | 0.0035 | -19.65 | 16.03 | -0.0011 | 0.00028 | -3.69 |
| **Pauli** | MN | 6.00E-05 | 0.0035 | 0.29 | 14.8 | 0.0071 | 0.00047 | 15.19 |
| **Pauli** | STH | -0.00017 | 0.0027 | -0.87 | 14.88 | 0.0087 | 0.00039 | 22.33 |

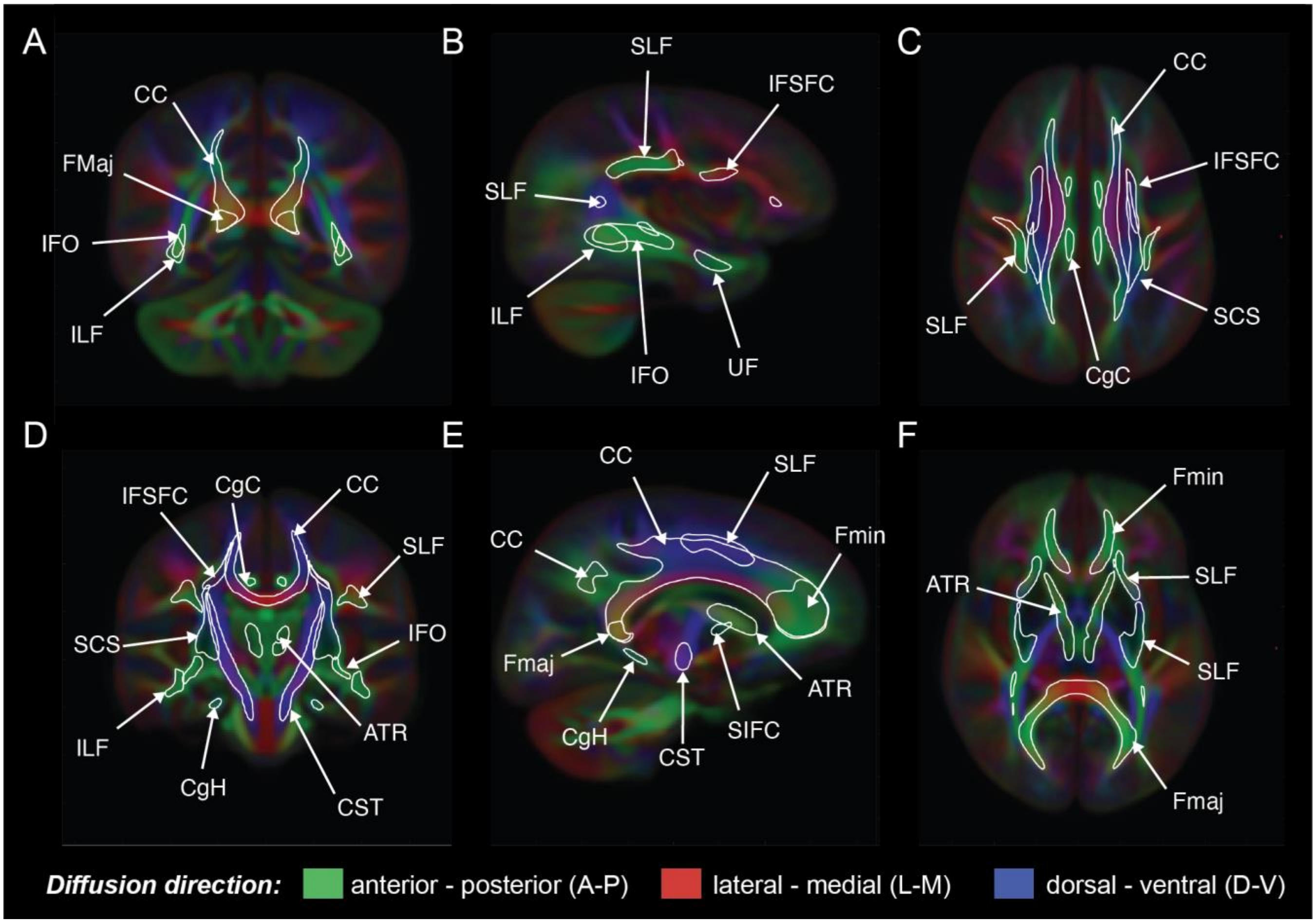

***Supplementary Figure 1. Voxelwise color-coded FA maps with labelled WM fiber tracts.*** *Mean voxelwise FA maps color coded based on the orientation of the primary diffusion direction: anterior-posterior (green), lateral-medial (red), and dorsal-ventral (blue). ROI outlines overlaid for the majority of the main WM fiber tracts generated using AtlasTrack (Hagler, et al., 2009). Brain slices (A-F) match those shown for Figures 1 and 2 in the main text. This provides a visualization of which tracts can be seen on the voxelwise statistical maps.*

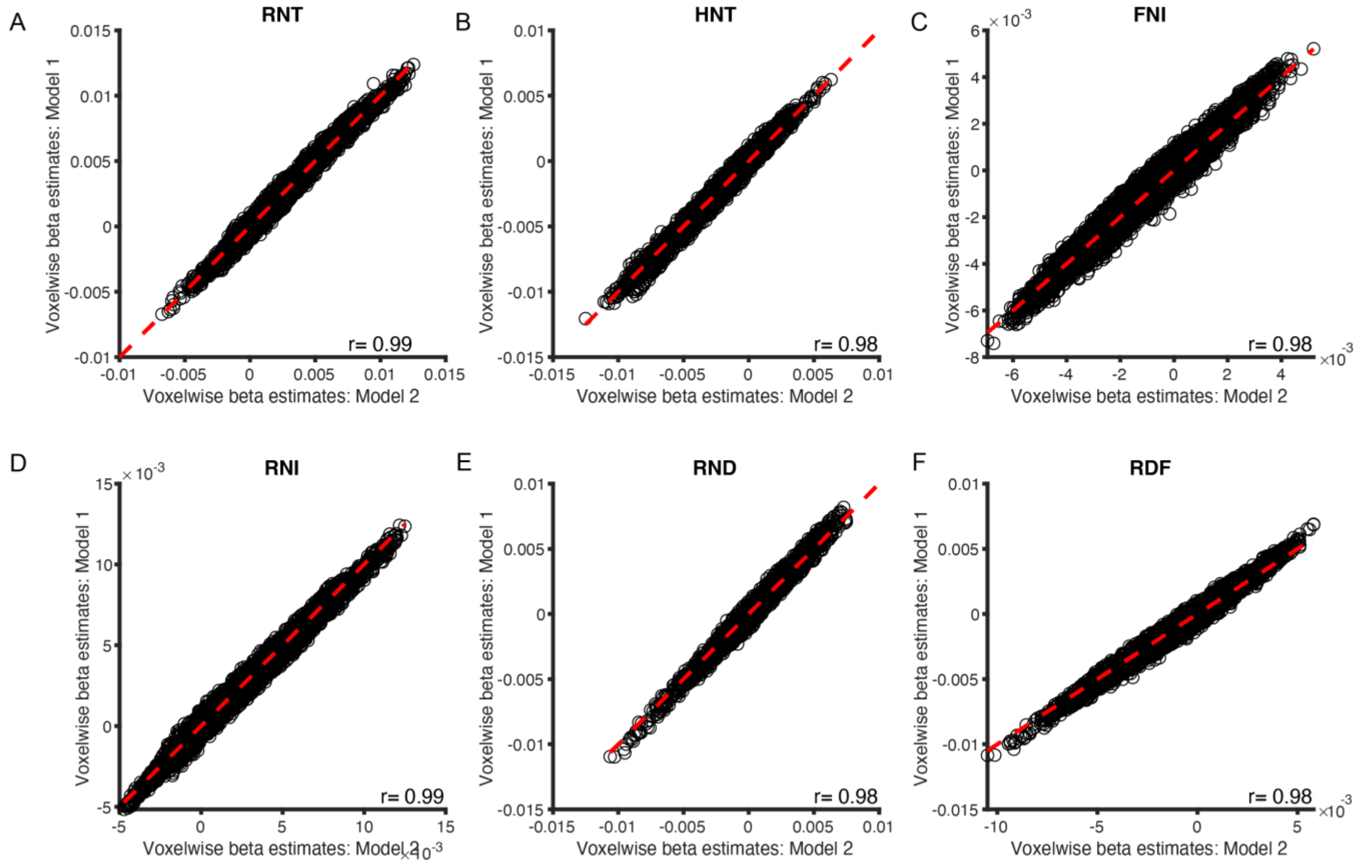

***Supplementary Figure 2. Correlations between voxelwise age associations with and without an age-by-sex interaction in the model.*** ***Bla****ck circles are voxelwise beta coefficients for the association between age and RNT (A), HNT (B), FNI (C), RNI (D), RND (E), RDF (F). Model 1 does not include the age-by-sex interaction (y-axis). Model 2 does include the age-by-sex interaction (x-axis). Red dotted line is a reference line with slope=1 and intercept=0. Pearson correlation coefficient is in the bottom right corner of each subplot.*

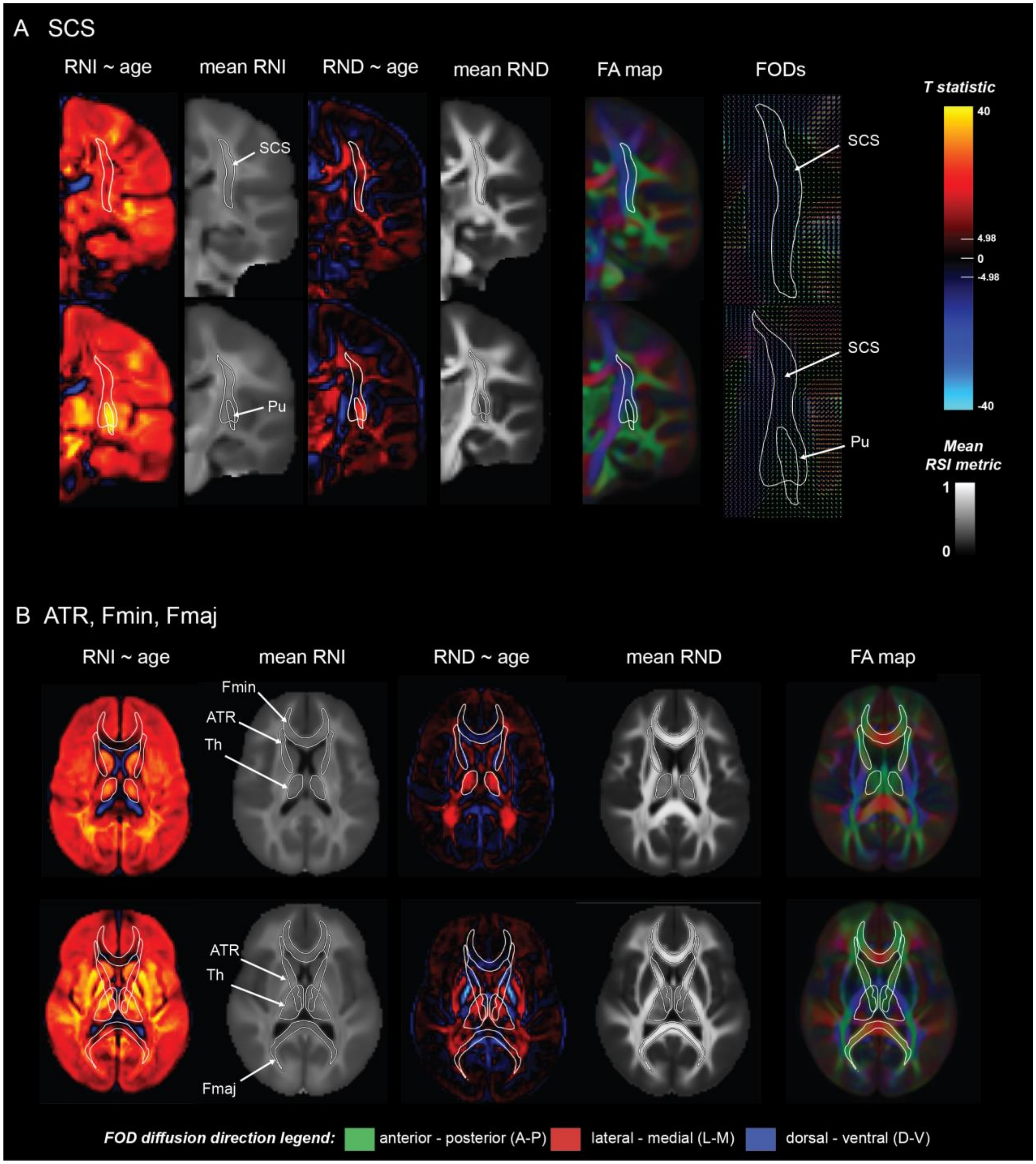

***Supplementary Figure 3. Associations between age, RNI and RND in specific tracts.*** *Unthresholded voxelwise t-statistics are shown for the SCS (A) and ATR (C) for RNI (column 1) and RND (column 3) alongside voxelwise mean RNI (column 2) and voxelwise mean RND (column 4) and color-coded FA maps (column 5). For panel A, a close-up of the voxelwise average FODs in the SCS and Pu (column 6). A) The SCS fiber tract outline is shown alongside the outline of the Pu in two slices moving from posterior (top image) to anterior (bottom image). Positive associations were greater and more significant in voxels overlapping with the Pu where the primary orientation of diffusion changes from dorsal-ventral (blue) in superior voxels of the SCS to anterior-posterior (green) in voxels in the Pu. B) Anterior portions of the ATR adjacent to the Fmin (top images) showed lower and less significant RNI and RND age associations compared to posterior inferior portions of the ATR (bottom images). Voxels in the ATR overlapping with the thalamus showed greater age-related changes in RNI and RND than other parts of the ATR. WM tract ROI abbreviations described in Supplementary Table 2. Effects are unthresholded. Voxelwise Bonferroni corrected significance threshold (|t|=4.98) is marked on the colorbar.*

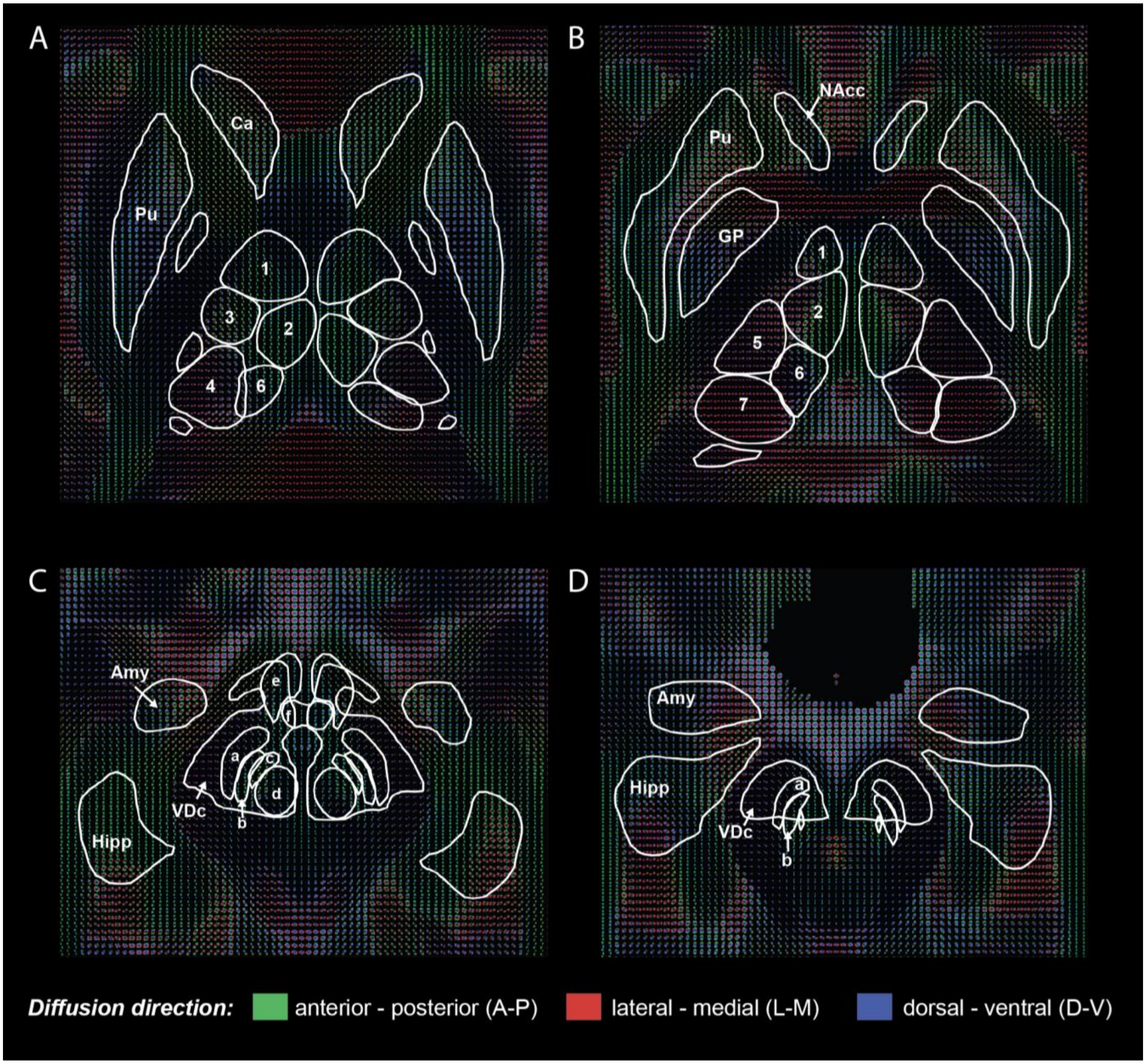

***Supplementary Figure 4. Voxelwise FODs averaged over participants within subcortical regions.*** *Voxelwise FODs, averaged across participants, show the orientation structure of diffusion in each voxel and are colored based on the diffusion direction (green=anterior-posterior; red=lateral-medial; blue=dorsal-ventral). There was clear variability in the orientation structure of diffusion within gross subcortical ROIs and the surrounding WM highlighting the importance of voxelwise analyses. By including atlases with a finer subcortical parcellation, we were able to localize age-related effects within large subcortical structures. ROI outlines here are from three different atlases. FreeSurfer aseg ROIs: Ca, Pu, GP, VDc, Amy, Hipp. Pauli atlas ROIs shown are: SNpc (a); SNr (b); PBP (c); RN (d); Hyp (e); MN (f). Najdenovska thalamic nuclei ROIs shown are: A (1); MD (2); VA (3); VLD (4); VLV (5); CP (6); P (7). ROI abbreviations are described in supplementary table 3.*

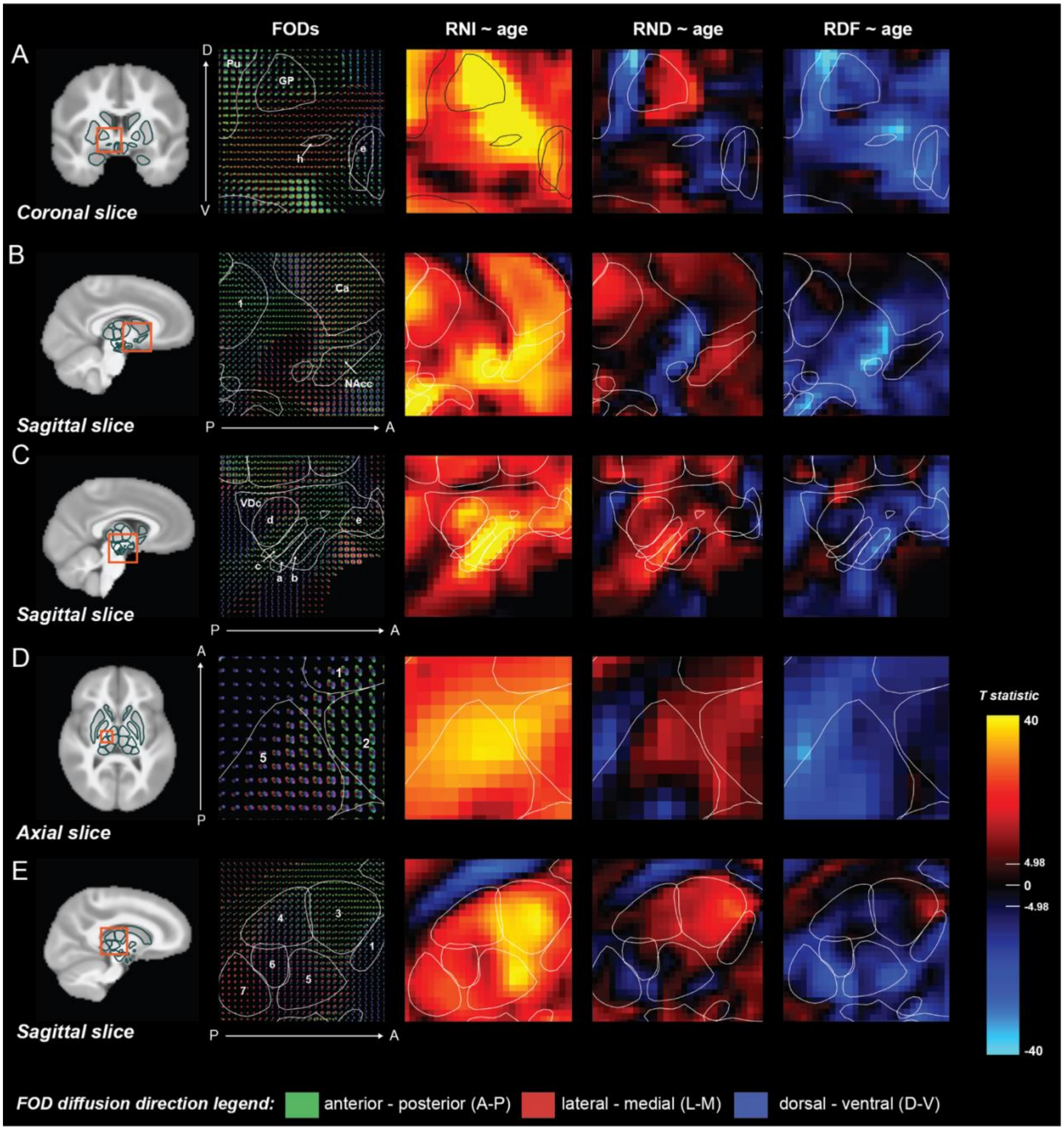

***Supplementary Figure 5. Zoomed in images of the voxelwise age associations with RNI, RND and RDF and mean FODs across different brain slices***. *A-D) Column 1: T1 slice showing orientation in the brain with orange box outlining the zoomed in area; column 2: mean FODs across participants; columns 3-5: voxelwise t-statistics for RNI, RND and RDF respectively. A) Coronal view of the GP, VP(h), and Hyp(e) showing highly significant RNI effects extending from the GP into the ventral region encompassing the VP and anterior commissure with primarily lateral-medial (L-M; red) diffusion; B) Sagittal view showing associations within the Ca extending into the ventral striatum and Hyp with both L-M and anterior-posterior (A-P; green) diffusion; C) Sagittal view of the VDC with ROIs of the RN (d), PBP (c), SNpc (a), SNpr (b) and Hyp (e) overlaid. The largest RNI and RND associations here can be seen in the SNpc, SNpr and PBP extending into the surrounding WM along voxels with diffusion primarily in the A-P and dorsal-ventral (D-V, blue) direction. D) Axial view of the thalamus showing the intersection of the A(1), MD(2) and VLV(5) nuclei of the thalamus with diffusion occurring in perpendicular directions. Both RNI and RND show highly significant age associations at this intersection. E) Sagittal view of the thalamic nuclei showing the most significant age associations for RNI and RND in the VA (3) nucleus extending into the VLV (5) nucleus where diffusion is primarily in the anterior-posterior (A-P, green) direction. Many voxels within the central(6) and pulvinar (7) nuclei where diffusion was primarily in the L-M direction were showed no significant age association with RND. Effects are unthresholded. Voxelwise Bonferroni corrected significance threshold (|t|=4.98) is marked on the colorbar.*

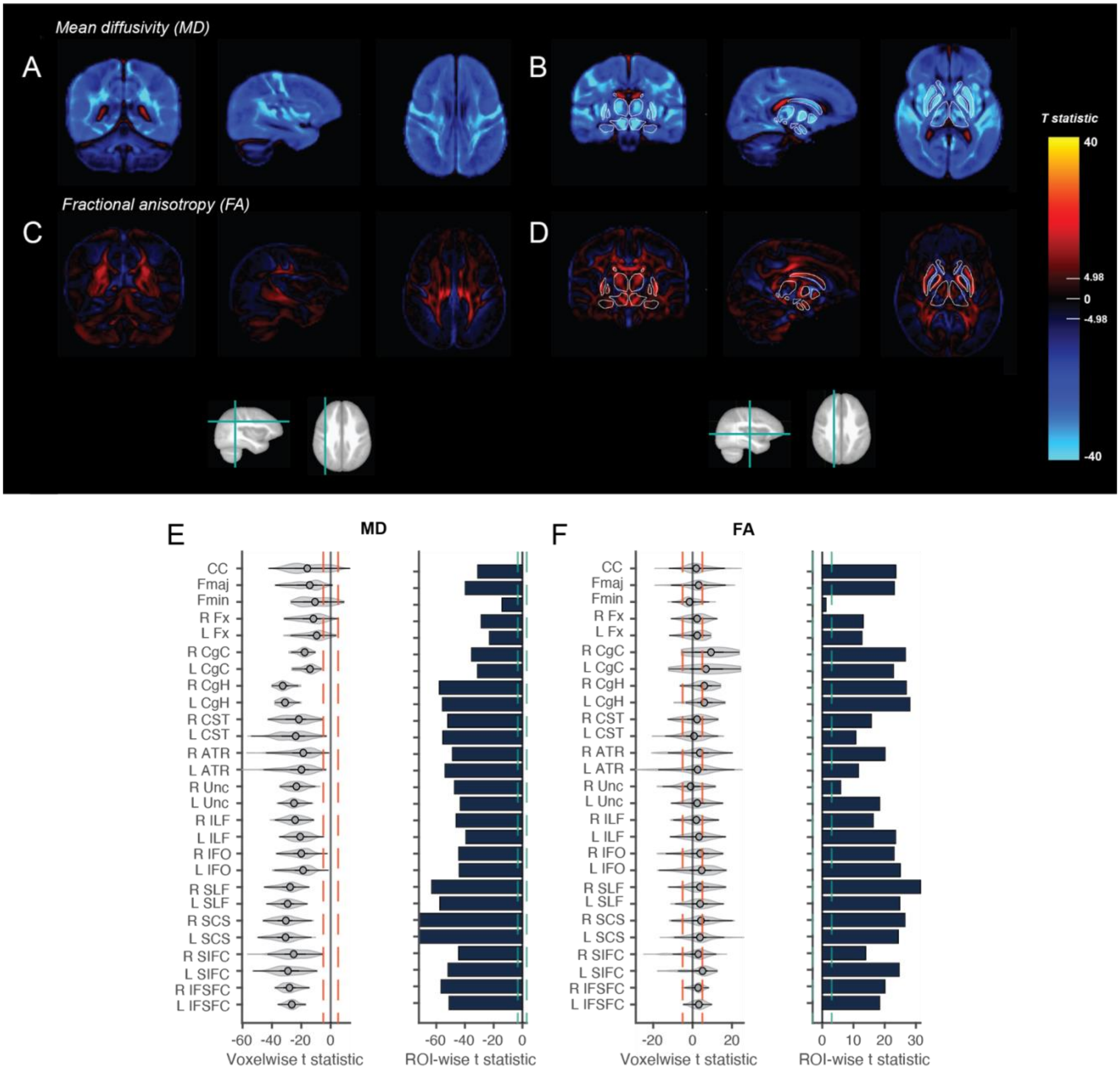

***Supplementary Figure 6. Associations between age and DTI metrics across the brain.*** *Voxelwise t-statistics for the association between age and MD (A,B) and FA (C,D) across different brain slices. Effects are unthresholded. Voxelwise Bonferroni corrected significance threshold (|t|=4.98) is marked on the colorbar. Outlines of the subcortical FreeSurfer ROIs are overlaid for the thalamus, caudate, pallidum, putamen, ventral diencephalon, amygdala and hippocampus to orient the reader. E-F) Violin plots show the distribution of voxelwise t-statistics extracted from each WM fiber tract. Red dotted lines show voxelwise Bonferroni corrected significance threshold. Bar plots show t-statistics from ROI analyses for the mean RSI metrics from each WM fiber tract. Green dotted line shows ROI Bonferroni corrected significance threshold (|t|=3.08). Plots are shown for MD (E) and FA (F). WM tract ROI abbreviations are outlined in Supplementary Table 2. Color coded FA in the same brain slices with WM fiber tracts labeled are shown in supplementary figure 1.*

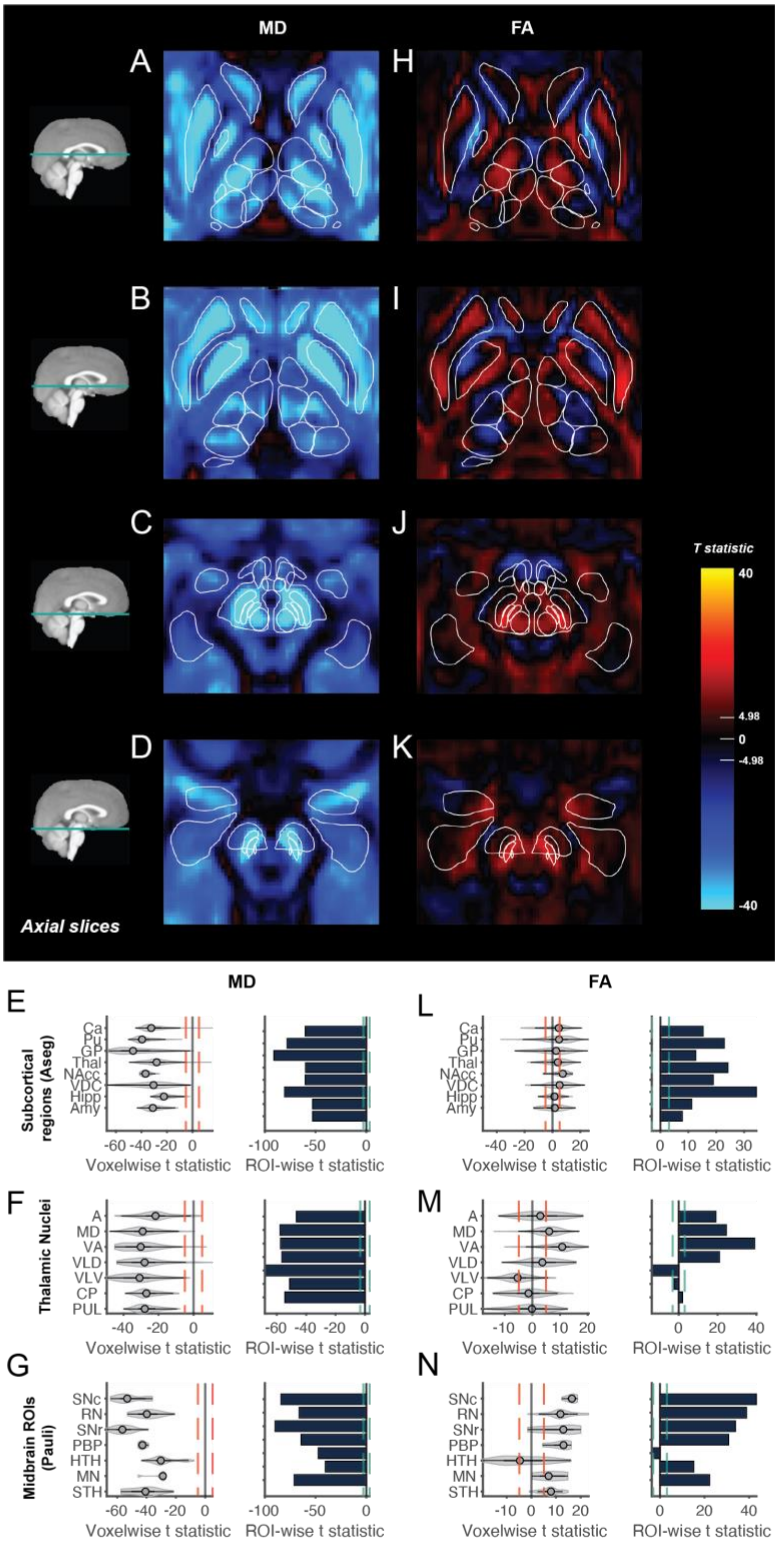

***Supplementary Figure 7. Associations between age and DTI metrics within subcortical regions.*** *Voxelwise t-statistics for the association between age and MD (A-D) and FA (H-K) across different axial brain slices moving from superior (top) to inferior (bottom). Effects are unthresholded. Voxelwise Bonferroni corrected significance threshold (|t|=4.98) is marked on the colorbar. Outlines of the Aseg, Pauli and Najdenovska ROIs are overlaid. Violin plots show the distribution of voxelwise age associations in each ROI for each RSI metric. Red dotted lines show voxelwise Bonferroni corrected significance threshold. Bar plots show t-statistics from ROI analyses for the mean RSI metrics from each subcortical ROI. Green dotted line shows ROI Bonferroni corrected significance threshold (|t|=3.08). Plots are shown for MD (E-G) and FA (L-N). Subcortical ROI abbreviations are outlined in Supplementary Table 3.*
